# Heritable temporal gene expression patterns correlate with metabolomic seed content in developing hexaploid oat seed

**DOI:** 10.1101/709774

**Authors:** Haixiao Hu, Juan J. Gutierrez-Gonzalez, Xinfang Liu, Trevor H. Yeats, David F. Garvin, Owen A. Hoekenga, Mark E. Sorrells, Michael A. Gore, Jean-Luc Jannink

**Affiliations:** Plant Breeding and Genetics Section, School of Integrative Plant Science, Cornell University, Ithaca, NY 14853, USA; Area de Genética, Departamento de Biología Molecular, Universidad de León, León, Spain; Corn Research Institute, Liaoning Academy of Agricultural Sciences, 84 Dongling Road, Shenyang 110161, China; USDA-ARS, Plant Science Research Unit, St. Paul, MN 55108, USA; Cayuga Genetics Consulting Group LLC,1430 Hanshaw Road, Ithaca, NY 14850 USA; USDA-ARS, Robert W. Holley Center for Agriculture and Health, Ithaca, NY 14853 USA

**Keywords:** transcriptome assembly, temporal gene expression, heritability, oat

## Abstract

Oat ranks sixth in world cereal production and has a higher content of health-promoting compounds compared to other cereals. However, there is neither a robust oat reference genome nor transcriptome. Using deeply sequenced full-length mRNA libraries of oat cultivar Ogle-C, a *de novo* high-quality and comprehensive oat seed transcriptome was assembled. With this reference transcriptome and QuantSeq 3’ mRNA sequencing, gene expression was quantified during seed development from 22 diverse lines across six time points. Transcript expression showed higher correlations between adjacent time points. Based on differentially expressed genes, we identified 22 major temporal co-expression (TCoE) patterns of gene expression and revealed enriched gene ontology biological processes. Within each TCoE set, highly correlated transcripts, putatively commonly affected by genetic background, were clustered, and termed genetic co-expression (GCoE) sets. 17 of the 22 TCoE sets had GCoE sets with median heritabilities higher than 0.50, and these heritability estimates were much higher than that estimated from permutation analysis, with no divergence observed in cluster sizes between permutation and non-permutation analyses. Linear regression between 634 metabolites from mature seeds and the PC1 score of each of the GCoE sets showed significantly lower p-values than permutation analysis. Temporal expression patterns of oat avenanthramides and lipid biosynthetic genes were concordant with previous studies of avenanthramide biosynthetic enzyme activity and lipid accumulation. This study expands our understanding of physiological processes that occur during oat seed maturation and provides plant breeders the means to change oat seed composition through targeted manipulation of key pathways.

## Introduction

Oat ranks sixth in world cereal production (USDA, 2019), and has a high content of health-promoting compounds in comparison to other cereals. Historically, oat was used primarily as animal feed (Hoffman, 2011), but recently it has been increasingly used as a human food because of health benefits associated with lipids, functional proteins, and dietary fibers such as β-glucan (Rasane *et al.*, 2013). Oat also produces unique phenolic compounds known as avenanthramides (Avns), which have been reported to modulate signaling pathways associated with cancer, diabetes, inflammation, and cardiovascular diseases (Tripathi *et al.*, 2018).

Despite worldwide production of this nutrient rich food, genomic studies in oats have lagged behind other cereal grains. A robust and comprehensively annotated oat reference genome is not yet available, and a limited number of oat transcriptome analyses have been published. Differential gene expression (DGE) analyses for salinity stress tolerance (Wu *et al.*, 2017) and responses under phosphorus deficit (Wang *et al.*, 2018) have been conducted in seedlings and roots, respectively. The first *de novo* seed transcriptome assembly was generated by Gutierrez-Gonzalez et al. (2013). However, this version of the transcriptome included only 412 of 1440 (28.6%, **Table S1**) complete BUSCO plant genes (Waterhouse *et al.*, 2018).

Investigation of the transcriptome through time is useful for understanding physiological processes occurring during seed maturation and for conducting genetic improvement. Much effort has been made to understand biological processes underlying observed temporal gene expression patterns, including transcriptome studies in maize (Li *et al.*, 2014; Yi *et al.*, 2019), wheat (Li *et al.*, 2018; Wan *et al.*, 2008) and barley (Zhang *et al.*, 2016). However, in each case, only one cultivar was examined, which may reflect genotype-dependent or genotype-specific results and thus, may have limitations for plant improvement. To date, no global/temporal gene expression studies of developing seed have been conducted in oat.

Analysis of Avns and lipid biosynthetic genes through time can facilitate an understanding their metabolism. Three genes encoding 4-coumaroyl-CoA 3-hydroxylase (*CCoA3H*), caffeoyl-CoA 3-O-methyltransferase (*CCoAOMT*) and hydroxyanthranilate hydroxycinnamoyltransferase (*HHT*) were cloned by Yang *et al.* (2004), and are key genes involved in Avns biosynthesis in oat (Collins, 2011). Oat grain has higher oil content than wheat or barley (Banaś *et al.*, 2007; Liu, 2011), and, unlike other cereals, the majority of oat lipids (86–90%) are found in the endosperm (Banaś *et al.*, 2007). However, oat lipid biosynthetic genes have yet to be cloned, and neither Avn nor lipid biosynthetic gene profiles have been investigated.

High throughput sequencing, *de novo* transcriptome assembly and quantification technologies are continually improving (Grabherr *et al.*, 2013; Patro *et al.*, 2017) making it possible to quantify transcript expression with high precision in non-model species, even when a reference genome sequence is not available. Furthermore, the 3’ mRNA sequencing technology enables the generation of gene expression profiling data for hundreds of samples with high precision and reasonable cost (Kremling *et al.*, 2018; Moll *et al.*, 2014; Tzfadia *et al.*, 2018). Here, we generated full-length transcript RNA sequences for developing seed of the oat cultivar Ogle-C (cv. Ogle-C) using both Illumina HiSeq 2000 and MiSeq sequencing platforms, together with QuantSeq 3’ mRNA sequencing data of developing seeds from 22 oat cultivars in two environments across six developmental time points. Our objectives were to: (i) generate a high-quality and comprehensive *de novo* oat seed transcriptome; (ii) identify global temporal gene expression patterns and reveal biological processes behind them; (iii) estimate heritabilities of identified temporal gene expression^1^ sets and evaluate their potential usefulness in plant breeding; and (iv) describe the temporal expression patterns of Avns and lipid biosynthetic genes.

## Results

### Validating the assembled oat transcriptome

The set of longest isoforms from each Trinity “gene” consisted of 134,418 transcripts (**Figure 1**). We aligned the Trinity longest isoform set against the *Brachypodium distachyon* (UP000008810), *Hordeum vulgare* (UP000011116), and *Triticum aestivum* (UP000019116) predicted proteomes (Uniprot 2019) using NCBI blastx (Camacho *et al.*, 2009), retaining 48,740 (36.26%) transcripts with at least one hit with an E-value < 10^−10^. The remaining 85,678 transcripts were aligned to scaffolds of the hexaploid oat genome v1.0 (*Avena sativa* v1.0, http://avenagenome.org/) using GMAP (Wu and Watanabe, 2005); 71,982 (53.55%) transcripts aligned with > 85% identity and > 85% coverage and 13,696 (10.19%) not aligning. The unaligned transcripts were queried using NCBI blastx against UniRef100 at E-value < 10^−3^. 3,879 transcripts were found to have at least one match, with 918 transcripts mapping to Viridiplantae (green plant) proteins and the remaining 2,961 transcripts mapped to non-Viridiplantae proteins. The 2,961 transcripts were excluded in the downstream analysis due to likely contamination. Therefore, our final representative transcriptome assembly (RTA) contained 131,457 transcripts (**Appendix S1**).

**Figure 1.**
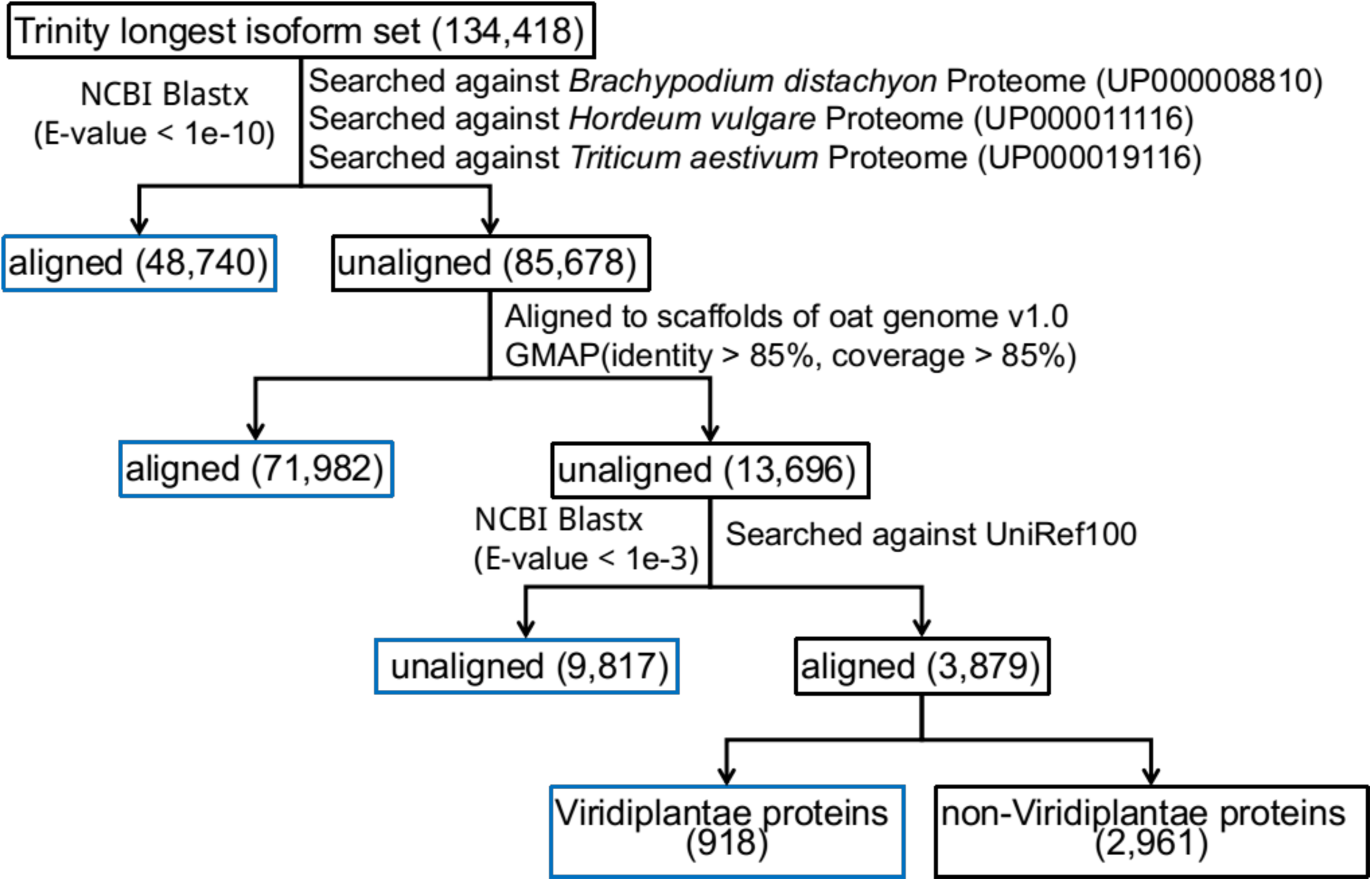
Results of aligning the assembled oat seed transcriptome against reference proteomes of oat relatives, scaffolds of the hexaploid oat genome v1.0, and the UniRef100.

56,877 (42.27%) and 27,278 (20.75%) transcripts of the RTA were longer than 500 and 1000 nucleotides (nt), respectively (**Table 1, Figure S1**). The N50, median and average transcript lengths were 1205, 433 and 757 nt, respectively. The RTA sums to a 99,539,633 nt assembly length.

**Table 1.** Statistics of transcriptome assembly and BUSCOs plants set assessment.

|  |  |
| --- | --- |
| Transcriptome assembly statistics |  |
| Total transcripts | 131,457 |
| Transcripts ($\geq 500$ nt) | 56,877 |
| Transcripts ($\geq 1000$ nt) | 27,278 |
| Contig N50 (nt) | 1,205 |
| Median contig length (nt) | 433 |
| Average contig length (nt) | 757 |
| Total assembled bases (nt) | 99,539,633 |
| BUSCO Statistics | number of genes (%) |
| Complete BUSCOs | 1212 (84.2%) |
| Complete and single-copy BUSCOs | 1188 (82.5%) |
| Complete and duplicated BUSCOs | 24 (1.7%) |
| Fragmented BUSCOs | 148 (10.3%) |
| Missing BUSCOs | 80 (5.5%) |
| Total BUSCO groups searched | 1440 |

nt = nucleotides, PE = paired-end

We used BUSCO (Waterhouse *et al.*, 2018) database to validate representation of protein-coding sequences in the RTA. Using the BUSCO plants set (embryophyta_odb9), 1212 of the 1440 BUSCO genes were complete in the RTA with 1188 genes single-copy and 24 duplicated (**Table S1**); 148 BUSCO genes were fragmented and 80 were missing (10.3% and 5.5% of the total, respectively).

### Principal components analysis (PCA) of samples

Developing seeds of 22 cultivars (**Table S2**) were collected at 8, 13, 18, 23, 28 and 33 days after anthesis (DAA) and expression abundances determined using 3’ mRNA Quant-Seq (Moll *et al.*, 2014). Of the 528 potential samples (22 lines × 6 time points × 2 sites × 2 replicates), we successfully sampled 419. From these 419 samples, 22 with less than 0.5 million mapped reads and 71,642 (53.30%) transcripts with less than two mapped reads in at least ten samples were removed. After this filtering, 397 samples (59,815 transcripts) were retained (**Appendix S2**). We performed PCA based on 500 transcripts with the highest variance. The second principal component separated the 397 samples into two distinct clusters with 71 and 326 samples (**Figure S2**). We could not identify the cause of this clustering. The PCA of the 326 samples showed the first principal component, explaining 64% of the variance, was driven by sampling time (**Figure 2**). Within the 326 samples, the average Pearson correlation coefficients of biological replicates were 0.874, 0.884 and 0.875 from Greenhouse samples, Field samples and among samples across the sites (**Figure S3**), respectively.

**Figure 2.**
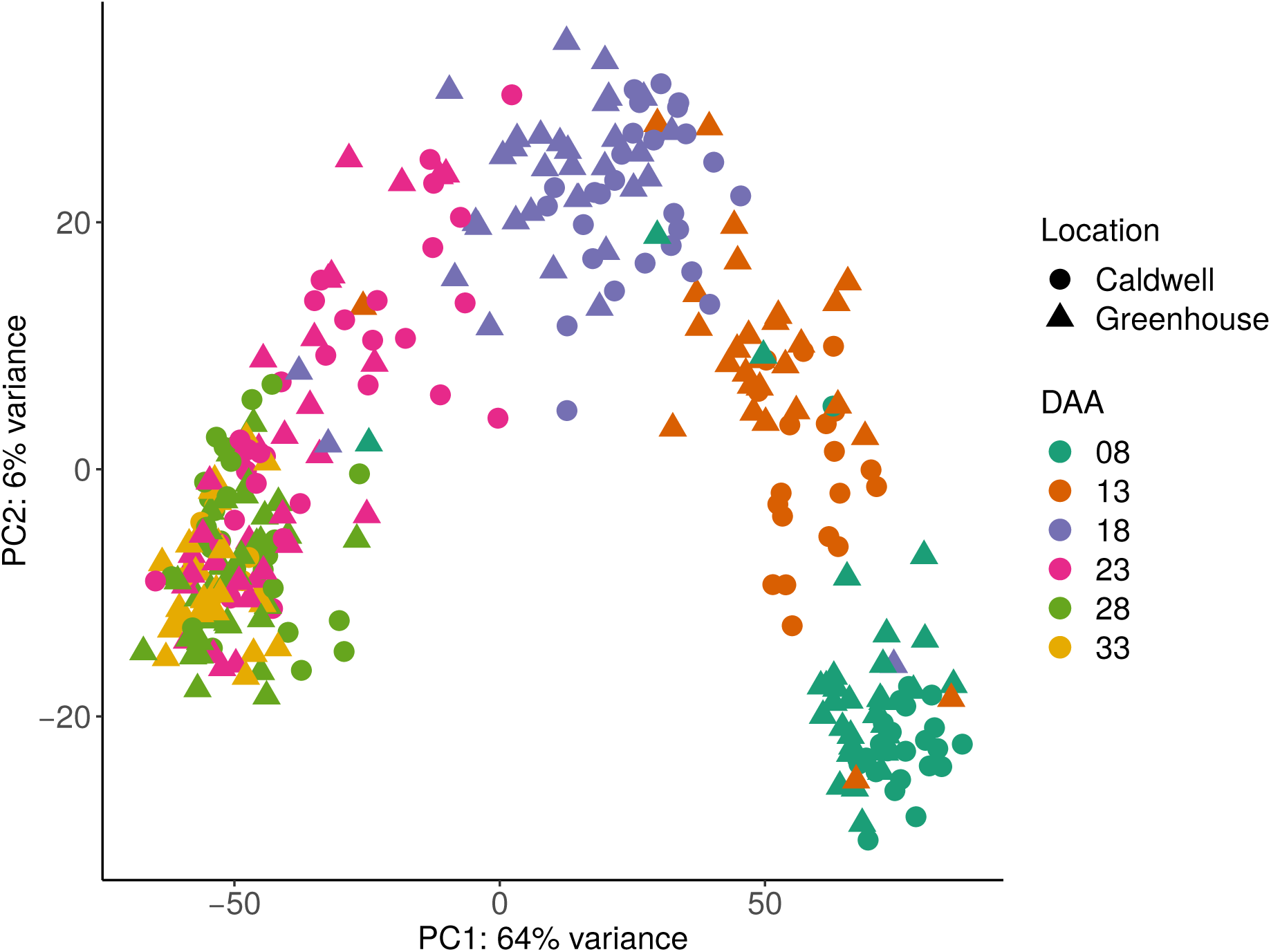
PCA plot of 326 samples with more than 0.5 million mapped reads based on the 500 transcripts with highest variance.

### Pairwise correlation of transcript expression and number of differentially expressed transcripts (DETs) between adjacent time points

Using the 326 samples, we performed pairwise correlation analysis and differential gene expression analysis between time points (**Appendix S3**). This analysis showed high correlation between adjacent time points, with decreasing correlation as time increased (**Figure 3a**). For example, the transcriptome expression of 8DAA had correlation coefficients of 0.959, 0.917, 0.817, 0.781, and 0.777 at 13DAA, 18DAA, 23DAA, 28DAA and 33DAA, respectively. This analysis also split the six time points into two groups. Transcriptome expression at 8DAA, 13DAA, and 18DAA showed higher correlation with each other than with later time points. Likewise, expression at 23DAA, 28DAA, and 33DAA showed higher correlation than with earlier time points. The lowest correlation between adjacent time points happened between 18DAA and 23DAA.

**Figure 3.**
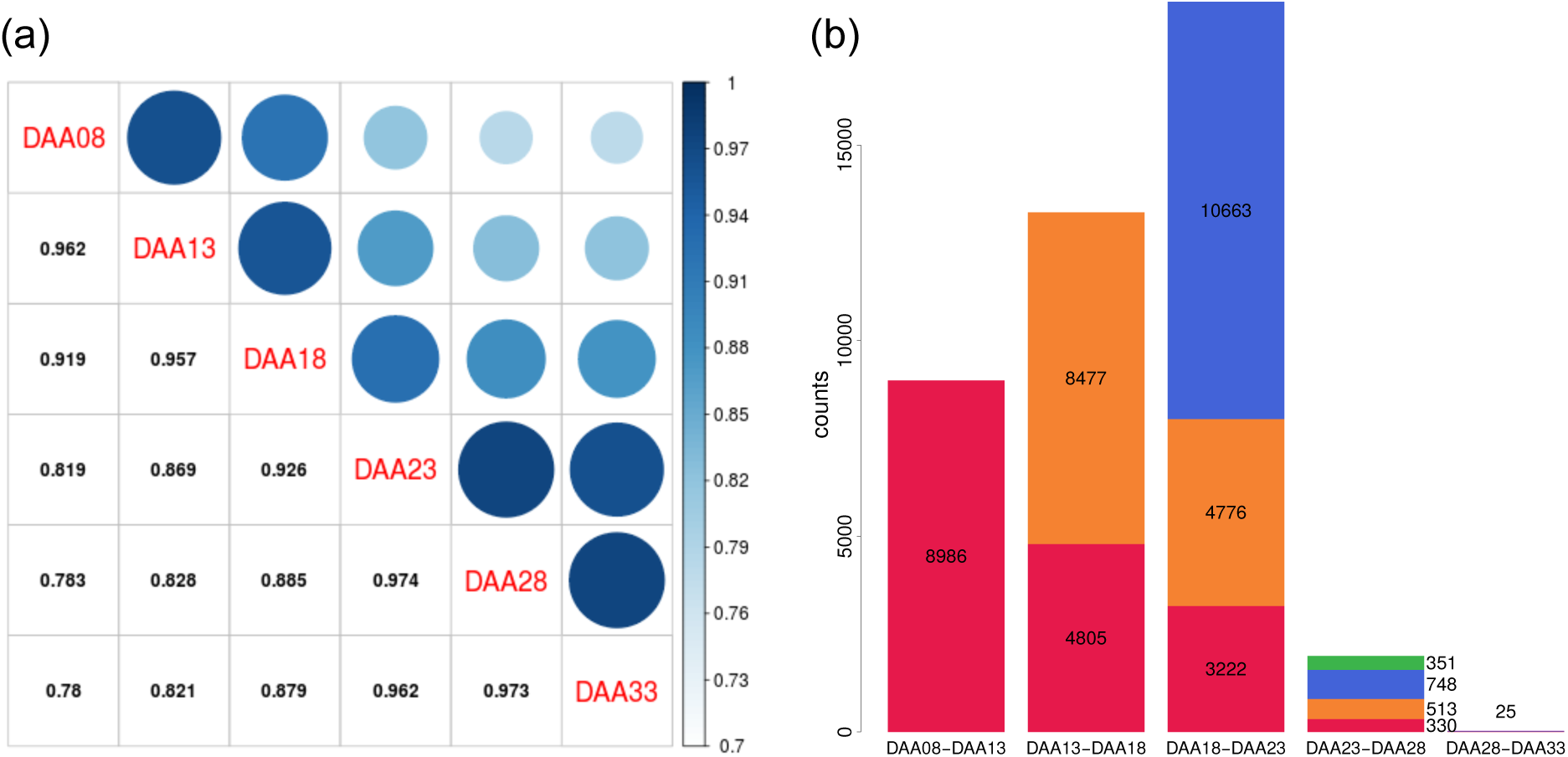
Pairwise correlation of transcript expression between time points (a) and numbers of differentially expressed transcripts between adjacent time points (b). In (b), total stacked bar height indicates the number of transcripts whose transcription level changed significantly over the time interval. Colored stack components indicate transcripts with significant change in an interval in common with a previous interval. For example, in red, 4805 transcripts significantly changed expression over both the 13-18 DAA and 8-13 DAA intervals.

DETs analysis between adjacent time points showed that the greatest number of DETs occurred between 18-23 DAA, and lowest number of DETs occurred between 28-33 DAA (**Figure 3b**). The maximum DETs occurred between early and middle stages of development, with many fewer DETs observed at later stages. We observed 8986 DETs between 8-13DAA, of which 4805 were also differentially expressed between 13-18DAA, while 8477 distinct transcripts were differentially expressed between 13-18DAA.

### Gene category (GO) over-representation analysis for DETs between adjacent time points

For DETs identified in each time interval, GO enrichment analysis (**Appendix S4**) was performed with all transcripts having at least one GO term as background set (Young *et al.*, 2010). The time interval of 13-18DAA had the highest number of over-represented GO categories at a false-discovery rate (FDR) adjusted p-value of 0.01 across all three domains of biological process, cellular compartments and molecular function (**Figure S4**), followed by 8-13DAA and 18-23DAA. Few GO categories were over-represented at 23-28DAA and 28-33DAA.

Generally, different GO categories were enriched for different time intervals, indicating the changing landscape of underlying processes. The common over-represented GO categories found between time intervals 8-13DAA and 13-18DAA were mainly related to peptide biosynthesis, amide biosynthesis and translation (**Figure S4a**). The common over-represented GO terms between time intervals of 13-18DAA and 18-23DAA related mainly to photosynthesis (**Figure S4a**,**c**). Oxidation-reduction (GO:0055114) was over-represented in all time intervals except 28-33DAA (**Figure S4a**), which is the very end of the sampled seed development stage. In contrast, nutrient reservoir activity(GO:0045735) was over-represented only in 28-33DAA (**Figure S4c**).

### Global temporal co-expression(TCoE) patterns

We used 25,971 total DETs between five pairs of adjacent time points to explore global TCoE patterns. Transcripts were clustered according to differential expression patterns. In theory, there are 3^5^=243 possible expression patterns considering that there are three states (up-regulated, down-regulated, and no change) in each of the five time intervals. We observed only 80 expression patterns (**Figure S5**) with a very skewed frequency such that the top 20 patterns contain 91% of the transcripts (**Figure 4**). A permutation test including 1000 permutations to simulate the null hypothesis that expression change in one time period is independent of that in other time periods showed a minimum number of 91 expression patterns compared to the observed 80 patterns, and a maximum of 84% of transcripts in the top 20 patterns compared to the observed 91% of transcripts. Relative to the permutation test, we observed far more transcripts whose expression changed in only one direction (either only going up or only going down over time) than would be expected: 79.7% of observed DETs change in only one direction compared to an null-hypothesis expectation of 50.0%. Among transcripts whose expression did reverse directions, we observe fewer transcripts going down then up (41.8%) than expected (45.7%). Both deviations were beyond the maxima from 1,000 permutations of the null hypothesis. Changes of state that included 13DAA or 18DAA were associated with the 6 largest (15249, 58.72% DETs associated) patterns, where one-step-up-at-18DAA (n = 5,006) and one-step-down-at-18DAA (n = 2,885) were the largest. Interestingly, we found transcript numbers in symmetrical expression patterns to be similar. For example, the expression pattern of one-step-up-at-8DAA (Top-8) is symmetrical to one-step-down-at-8DAA (Top-10), and they contain a comparable number of transcripts (928 and 878, respectively). We also found that the number of transcripts in an expression pattern was predicted by the number of differential expression events in the pattern (e.g., “One-step-up-at-8DAA (Top-8)”, “Up-at-13DAA-down-at-18DAA (Top-19)”, and “Three-steps-up-at-8DAA (Top-9)” have one, two, and three differential expression events, respectively. **Figure S6**). Thus, the number of transcripts in an up-regulated pattern was well explained by the number of transcripts in its symmetrical down-regulated pattern and by its number of differential expression events.

**Figure 4.**
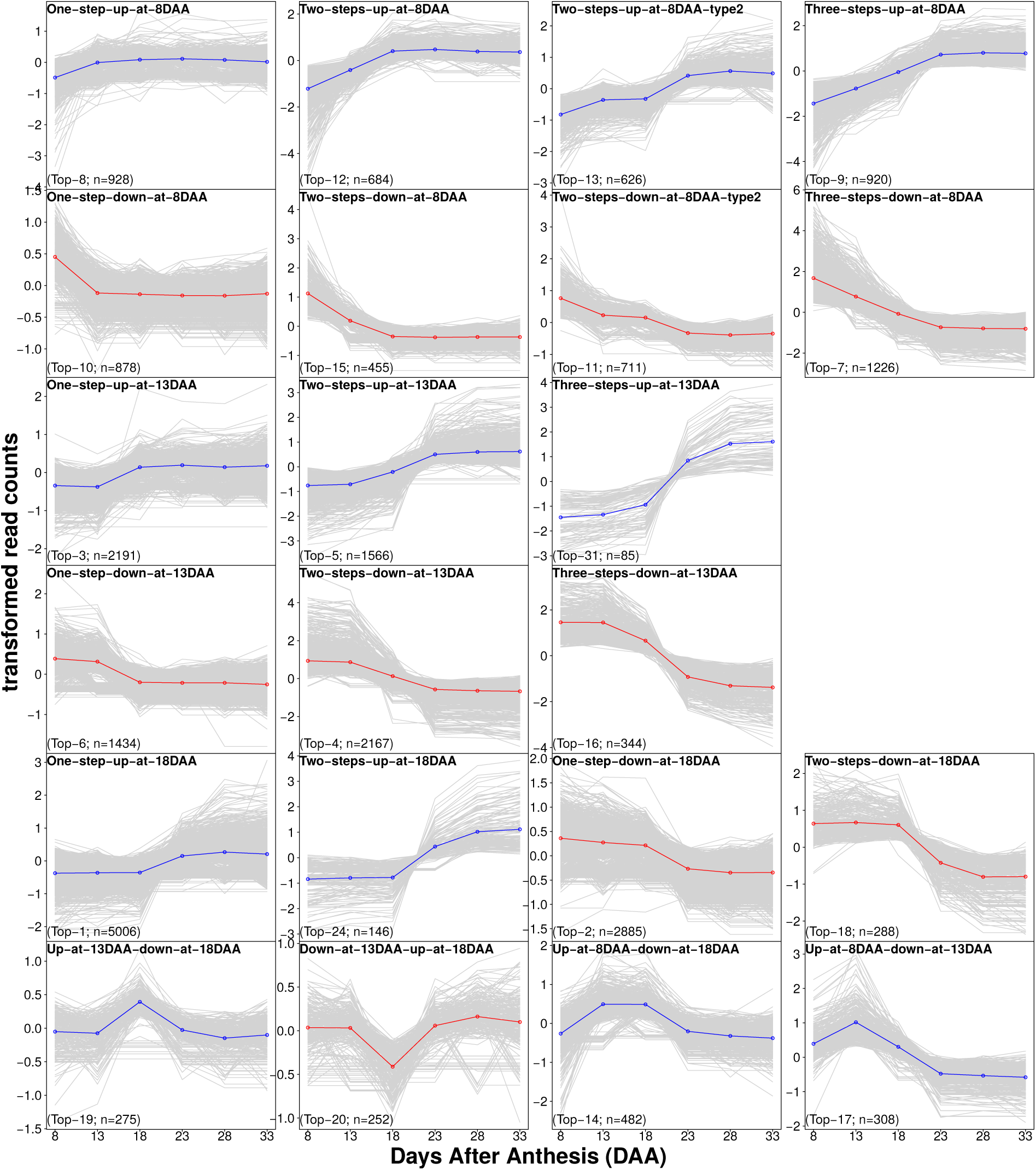
Top20 expression patterns plus two main expression patterns of 13DAA (Three-steps-up-at-13DAA, Top-31) and 18DAA (Two-steps-up-at-18DAA, Top-24). Expression pattern plots are named, and ranking numbers and total number of transcripts of each expression patterns are given at the bottom. Median gene expression profiles of individual transcripts across the 22 oat lines were depicted in gray lines, and average expression profiles for each pattern are depicted in blue (if up-regulated) or red (if down-regulated).

### Gene ontology analysis for identified TCoE sets

For the groups of genes identified by temporal clustering, 8 of the 22 patterns exhibited significant GO enrichment (**Figure S7**). The One-step-down-at-8DAA (Top-10) pattern exhibited GO enrichment for tRNA aminoacylation (protein translation), amino acid activation and tRNA aminoacylation. Two-steps-down-at-8DAA (Top-15) was associated with cellular localization and intracellular protein transport processes. GO terms enriched for the Three-steps-down-at-8DAA (Top-7) included nucleosome assembly and protein-DNA complex assembly related processes. GO terms enriched for the One-step-up-at-13DAA (Top-3) included a group of complex processes related to peptide biosynthesis, translation, rRNA and ncRNA processing/metabolic and nucleic acid metabolic processes. GO terms enriched for the Two-steps-up-at-13DAA (Top-5) included a group of processing/metabolic procedures related to rRNA, ncRNA, mRNA and tRNA. GO terms enriched for the Two-steps-down-at-13DAA (Top-4) included a group of processes related to photosynthesis. GO terms enriched for the One-step-up-at-18DAA (Top-1) included a group of processes related to regulation of biological process, gene expression, cellular process, metabolic process etc. GO terms enriched for the Up-at-step-8DAA-down-at-13DAA (Top-17) related to regulation of photosynthesis.

### Heritability estimation of the 22 TCoE sets

As we had both temporal and genetic breadth in our design, we estimated the heritability of our transcriptome, a novelty compared to previous studies (Li *et al.*, 2014, 2018; Wan *et al.*, 2008; Yi *et al.*, 2019; Zhang *et al.*, 2016). Within a TCoE set, we asked two questions: (i) which transcripts are strongly correlated with each other across genotypes such that they are similarly affected by genetic background? And (ii) Is sufficient variation in transcript expression explained by genotype so that it potentially can be manipulated by plant breeders to change oat seed composition?

With these aims, we created subclusters (varying in number from 4 to 13) within each TCoE set based on the adjusted gene expression matrix of the full set of 397 samples with more than half million mapped reads (see Methods for details). We termed such subclusters genetic co-expression (GCoE) sets. Within each GCoE set, we calculated the PC1 score for each of the 22 oat lines, and estimated the additive genetic variance of that score. We applied the same procedure to 50 permuted data sets. Heritabilities estimated from GCoE sets of the 22 TCoE sets (real data) were much higher than those estimated from permuted datasets (**Figure 5**). A majority of TCoE sets had highly heritable GCoE sets. Fifteen TCoE sets had GCoE sets with median heritablity exceeding 0.75; two had median heritabilities of GCoE sets between 0.5 and 0.75; three had median heritabilities of GCoE sets between 0.25 and 0.50. Two TCoE sets had GCoE sets with median heritability less than 0.25.

**Figure 5.**
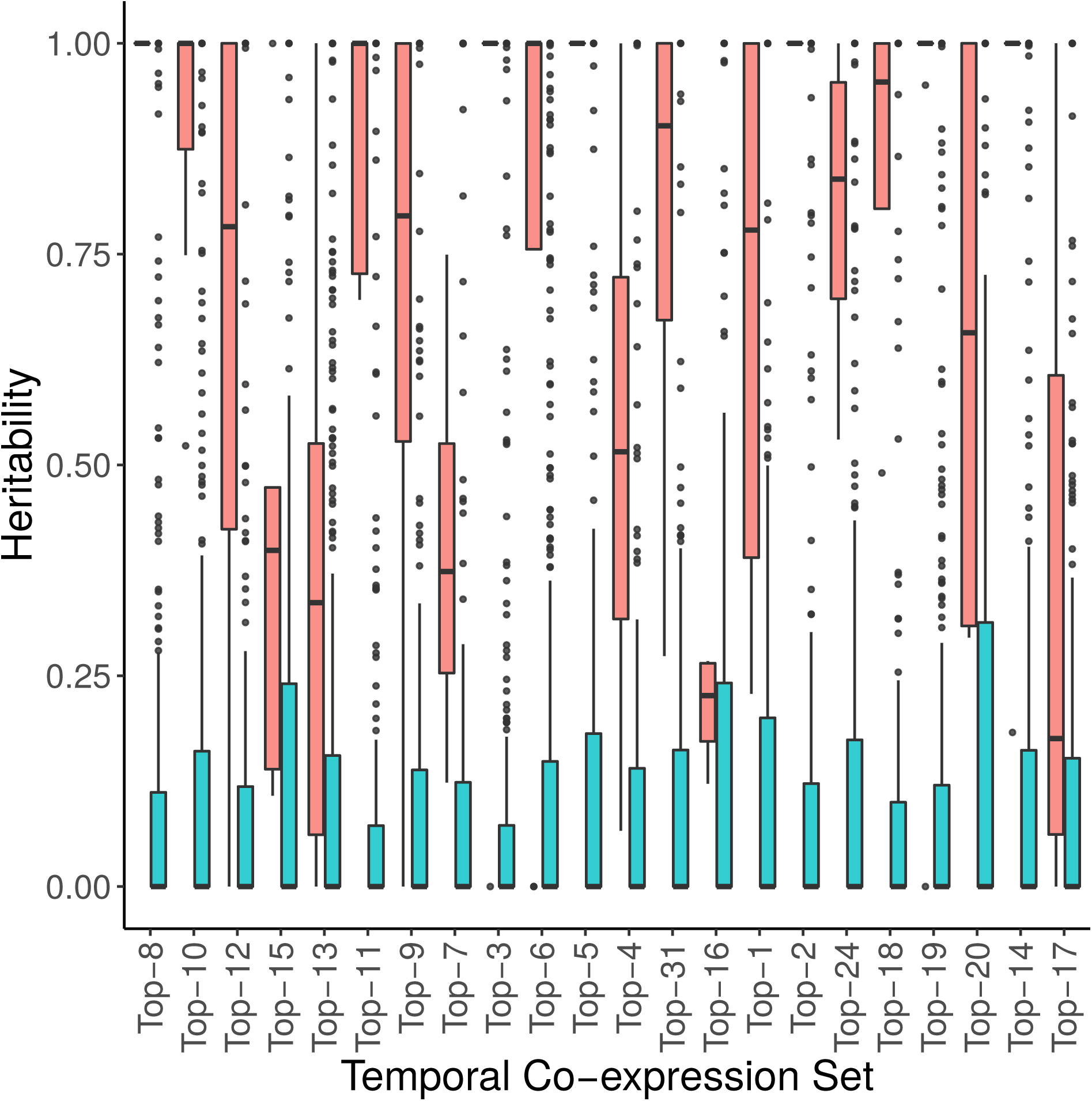
Box plots of heritabilities estimated from GCoE sets of the 22 TCoE sets (red) against box plots of heritabilities estimated from permuted TCoE sets (blue). The TCoE sets were ordered as in Figure 4.

To test whether the distribution of GCoE set sizes of the 22 TCoE sets differed from that expected under the null distribution, we performed permutation analyses. We calculated Mahalanobis distance of cluster sizes from 1,000 permutations to generate a Mahalanobis distance distribution of each permutation from the mean. We then calculated the Mahalanobis distance of the cluster size vector of the non-permuted expression matrix to the mean of permutation-based Mahalanobis distances, and tested it using a standard Chi-squared test, since the squared Mahalanobis distance follows a Chi-Square distribution (Brereton, 2015; Wicklin 2012). None of the 22 TCoE sets deviated from the null distribution constructed from permuted datasets at significant level of 0.05 after Bonferroni correction (**Table S3**).

### Correlation between transcript expression patterns and metabolites

To examine whether the transcript expression patterns associated with metabolite abundance, we applied a simple linear regression to detect the relationship between 634 metabolites (each with heritability > 0.4) of mature seeds and PC1 scores of GCoE sets. The 634 metabolites included 9 fatty acids, 199 and 426 metabolite features obtained from targeted GC-MS, non-targeted GC-496 MS, non-targeted LC-MS analyses, respectively. For almost all. We applied the same the p-values from real data were much lower than that obtained from permutations (**Figure S8**).

### Temporal transcript expression pattern of Avns and lipid biosynthetic genes

Two of the compositional features that distinguish oats from other cereals are high lipid levels and the multifunctional Avns. We identified transcripts with sequence similarity to biosynthetic genes for both pathways (**Table S4**). All of our candidates showed long alignment length and high percentage of identity to their reference sequences, and each had a high number of mapped reads across all samples except *FAD3*, which was excluded in expression pattern analysis.

*CCoA3H, CCoAOMT* and *HHT* are three key genes for Avns biosynthesis (Collins, 2011). The *CCoA3H* gene was up-regulated from 8DAA (**Figure 6a**), reaching a peak either at 13DAA or 18DAA depending on genotype and then declining, and reaching a plateau at 23DAA or 28DAA. The *CCoAOMT* gene showed a similar expression pattern to that of the *CCoA3H*, but with more variation between genotypes. Expression of the *HHT* gene moved up and down within a relatively small range across all time points, but did not show a clear expression pattern common across all genotypes.

**Figure 6.**
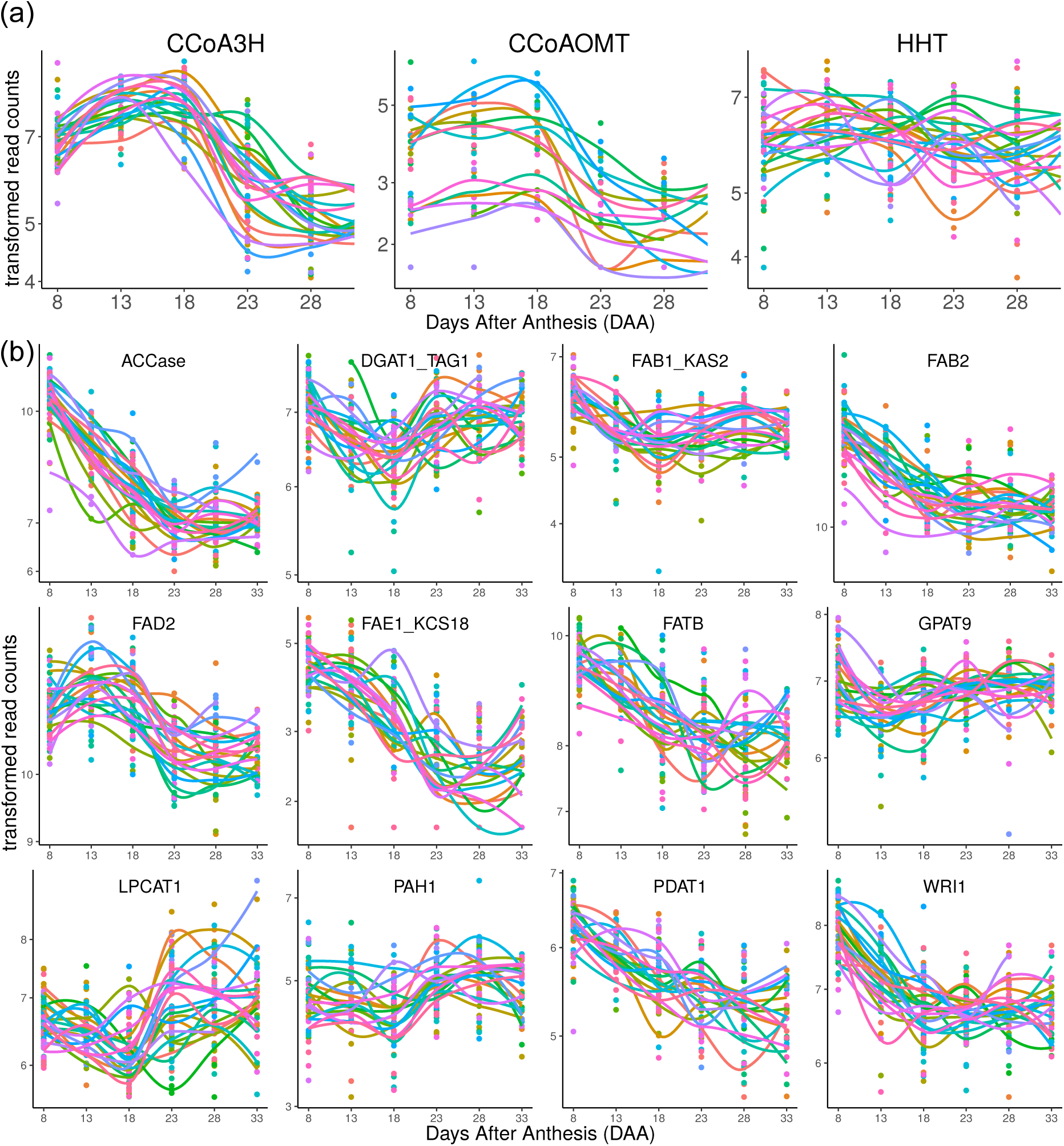
Transcript expression patterns of oat avenanthremides (a) and fatty acids (b) biosynthetic genes based on individual oat lines. Transcript expression values of individual samples were depicted in colored dots, and a LOWESS (Locally Weighted Scatterplot Smoothing) curve through all expression values of each genotype was drawn in colored smooth lines. Different colors represent different oat lines. Oat lines with too many missing values were excluded.

Key genes involved in fatty acid biosynthesis showed several different expression patterns (**Figure 6b**). Expressions of *ACCase, FAB2, FAE1/KCS18, FATB, PDAT1* and *WRI1* started to decline from 8DAA and reached a plateau either at 23DAA or 18DAA (*WRI1*). Expressions of *DGAT1*/*TAG1, FAB1*/*KAS2, LPCAT1* and *PAH1* started to decline at 8DAA, reached a valley at 18DAA, and rose to a plateau at 23DAA. Expression of the *FAD2* gene started to rise at 8DAA, reached a peak at 13DAA, and declined after 13DAA until reaching a final plateau at 23DAA. The *GPAT9* gene showed different expression patterns between genotypes, but most genotypes started to decline at 8DAA, then rose after 13DAA, and reached a final plateau at 23DAA.

## Discussion

### Transcriptome assembly validation and quality evaluation

A common issue for *de novo* transcriptome assembly is that while there are many transcripts in the initial assembly, there is no optimal approach to filter them. A number of studies have used the longest isoform (Gutierrez-Gonzalez, Tu, *et al.*, 2013; Hirsch *et al.*, 2014). In this study, we started with the longest isoform set (n = 134,418), and found 90.5% of it could be aligned to oat relatives (n=48,740), oat genome scaffolds (n=71,982) or Viridiplantae proteins (n=918, **Figure 1**). However, 9,817 (7.3%) transcripts couldn’t be aligned to any of these. Hypotheses to explain non-alignment are that they were too small to align to a protein in UniRef100 (**Figure S9**), were non-coding RNA, were sequence unique to oat, or Ogle-C specific transcripts missing in the oat genome v1.0. There was no good reason to filter them out, so we included them in our RTA for the downstream analyses.

Various methods have been proposed to assess the quality of transcriptome assemblies. BUSCO has been considered the gold standard to evaluate completeness of genome assembly for transcriptome assembly (Simão *et al.*, 2015). The BUSCO plant set (embryophyta_odb9) evaluates assembly content by searching the assemblies for 1440 conserved single copy orthologs found in at least 20 of 31 plant species (Waterhouse *et al.*, 2018). Of those, 1212 (84.2%) BUSCO plant genes were found to be complete in our RTA, which indicates a high level of overall coverage for our transcriptome assembly. Our dataset is a substantial improvement over the first oat seed transcriptome assembly (Gutierrez-Gonzalez, Tu, *et al.*, 2013), which only included 412 (28.7%) complete BUSCO plant genes (**Table S1**). Based on the expression profiles of 12 HiSeq samples of cv. Ogle-C whose developing seeds were collected at 7, 14, 21, and 28 DAA with three biological replications each, we were able to assign all 12 samples into four clusters corresponding to the four sampling times (**Figure S10**), and the average correlation among biological replicates was 0.97 (**Figure S11**). Finally, we evaluated the quality of our transcriptome assembly by searching the RTA for Avns and lipid biosynthetic genes homologous to other oat cultivars or other species. All three genes of *CCoA3H, CCoAOMT, HHT* involved in Avns biosynthetic pathways were found to have high similarity to their reference sequences from *Arabidopsis, Brachypodium distachyon* or other oat cultivars (**Table S4**). Twelve key genes involved in fatty acid biosynthesis were found to have high quality homologs in the RTA, with the alignment length ranging from 825 bp to 7598 bp and the percent identity ranging from 72.8% to 88.7%. For the *ACCase* gene, the *B. distachyon* reference sequence was 8783 bp, and the homologous transcript found in the RTA was 7812 bp with alignment length of 7598 bp and percent identity for the alignment region of 88.7%. In summary, we created a high quality and comprehensive transcriptome assembly, which is reliable for downstream analysis.

### Important biological processes underlie different oat seed development stages

In *Arabidopsis* seed development, major accumulation of storage proteins occurs between 5 and 13 days after flowering (Ruuska, 2002). In maize, Li et al. (2014) found that DGE in early seed development (0-10 DAA) related to storage protein preparation. In wheat grain development, Wan et al. (2008) identified storage protein transcripts most abundant at around 14 DAA. In our study, for the early stage of oat grain development (8-13 DAA and 13-18 DAA), the dominant biological process ontologies enriched included peptide biosynthesis, amide biosynthesis, organonitrogen compound biosynthesis and translation, which are all relevant to protein synthesis. This suggests that oat seed storage proteins also accumulate at early grain development stages between 8-18 DAA.

Li *et al.* (2014) observed rRNA and ncRNA related biological process ontologies enriched in early developing kernels of maize (0-10 DAA). We found rRNA and ncRNA related biological process ontologies enriched between 13-18 DAA, which indicates that rRNA and ncRNA processing procedures might also be important between 13-18 DAA in oat.

Expression of photosynthetic genes peaked at 11 days after flowering in *Arabidopsis* developing seeds (Ruuska, 2002). Photosynthesis is the dominant biological process ontology identified at 14 DAA in wheat grain development (Rangan *et al.*, 2017). Expression of 20 of 29 (68.97%) photosynthesis-related genes peaked at 8 DAA in developing barley grains (Bian *et al.*, 2019). Here, photosynthesis related GO terms were enriched in time intervals of 13-18 DAA and 18-23 DAA, which suggests immature oat seeds at early and middle development stages contain functional chloroplasts capable of photosynthesis during grain filling.

A GO category of nutrient reservoir activity was found enriched between 28-33 DAA, which suggested the importance of nutrient accumulation and storage at the late seed development stage. This GO term was also found to be enriched at storage phase of barley seed development (Bian *et al.*, 2019).

### Canalization and genetic differentiation of transcription

Given the 58,120 transcripts measured in the 3’ QuantSeq assay, we find it remarkable that only 494 showed a time by genotype interaction at an FDR of 0.1. While it is unclear how to formulate a null hypothesis against which to test this number, the fact that it is less than 1% of the transcripts suggests that temporal dynamics of expression are tightly controlled and canalized across genotypes. The seed is the sole vehicle for the survival of an annual from one year to the next. It stands to reason, therefore, that its composition, as affected by the temporal sequence of gene expression and therefore enzymatic activity is important to fitness. A characteristic the analysis revealed about seed gene expression is that it is unimodal: 92% of transcripts showing differential expression had only one peak of expression over the development of the seed. In other words, only 8% of transcripts showed first a significant drop followed by a significant rise in expression which would lead to expression peaks in distinct early and late periods of seed development. Only one of the top 22 clusters showed this pattern (Top-20 with 252 transcripts) and no gene ontology terms were enriched in this cluster.

While the temporal expression patterns appeared conscribed, our data also offered the possibility of exploring genetic variability in expression. To explore genetic variation, we further clustered transcripts in each TCoE set according to their co-expression across oat lines, allowing us to test the heritability of such genetic co-expression sets. We observed that for 17 of the 22 TCoE sets, median heritabilities of GCoE sets were above 0.50. Particularly, for 6 of the 22 temporal co-expression sets, heritabilities of GCoE sets were close to 1. The high heritabilities of GCoE sets arise for the following reasons: (i) within a GCoE set, profiles of transcripts are highly correlated and with almost the same shapes across genotypes, so the majority of variation in expression profiles is expected to be explained by variation of genotypes; (ii) PC1 was used to characterize a GCoE set, which reduced noise from individual transcript expression profiles (Krafft *et al.*, 2011).

We further found heritabilities of GCoE sets estimated from our real dataset were much higher than those estimated from permuted datasets (**Figure 5**). Moreover, after Bonferroni correction, none of the 22 temporal co-expression sets had a cluster size distribution significantly different from that of a null distribution obtained by permution (**Table S3**). Completing the causal chain from genotypes to transcribed genes to metabolomic phenotypes, we showed that for the overwhelming majority of GCoE (106 GCoE identified across 22 TCoE, **Figure S8**), transcript levels correlated with metabolite levels. Given the relatively small number of oat lines we worked with, statistical power to identify specific transcript to metabolite correlations was too low to overcome the multiple testing burden. Nevertheless, these correlations suggest the groups of genes we observed at temporally co-regulated clusters are biologically meaningful and represent useful groups of traits that breeders will able to select upon to manipulate oat seed composition to more desirable endpoints.

### Temporal transcript expression patterns of Avns and lipid biosynthetic genes

Avns are produced in both vegetative tissues and grain (Matsukawa *et al.*, 2000; Peterson and Dimberg, 2008; Wise, 2017). Enzymes involved in the biosynthetic pathway of the avenanthramides include *CCoA3H, CCoAOMT* and *HHT (Collins, 2011; Yang et al., 2004). HHT* is the final enzyme in the biosynthetic pathway. Little research has been conducted on gene expression of the three enzymes in oat. Activity of the final biosynthetic enzyme, *HHT*, has been found in dry seeds (Bryngelsson *et al.*, 2003; Matsukawa *et al.*, 2000). Temporal dynamics of *HHT* activity were investigated in spikelets containing developing grain using nine field-grown cultivars (Peterson and Dimberg, 2008). Most cultivars showed a trend of increasing activity during maturation, however, the *HHT* activity peaked at different times and had high variation at final harvest among cultivars (Peterson and Dimberg 2008). Similarly, in our study, we did not observe a clear common gene expression pattern across all 22 genotyes for the *HHT* gene, although both *CCoA3H* and *CCoAOMT* showed a similar expression pattern over maturation across most of cultivars. This might be attributed to the complex role the *HHT* enzyme plays in biosynthesis of Avns, as it is involved in three different pathways and catalyzes the biosynthesis of several different Avns (*Collins, 2011*).

In wheat and barley grains, oil accounts for 2–3% of seed dry weight (Barthole *et al.*, 2012). In contrast, oat grains are relatively rich in oil, which can vary from 3% to 11% of grain weight in different cultivars (Banaś *et al.*, 2007; Liu, 2011), with breeding lines containing up to 18.1% (Frey and Holland, 1989). In most cereal grains, oil is mostly stored in the form of triacylglycerols (TAGs, esters of fatty acids and glycerol) within the embryo. However, the majority of oat lipids (86–90%) are found in the endosperm, and up to 84% of the lipids are deposited during the first half of seed development, when seeds are still immature with a milky endosperm (Banaś et al. 2007). Little research has been done on temporal expression of genes related to oil storage in cereals. In the barley embryo, most lipids were deposited between 12-22 DAA, and the temporal expression profile of the *oleosin 2* transcript constantly increased between 8-22 DAA, and declined thereafter (Neuberger 2008). However, we observed most lipid synthesis genes had high expression level at 8 DAA, and then were down regulated, maintaining a low expression level after 23 DAA. This is distinct from barley lipid synthesis gene expression, but consistent with findings of Banaś *et al.* (2007) that most oat lipids were deposited at early and middle stages of seed development.

## Experimental procedures

### Sample collection, RNA extraction, cDNA construction, Illumina sequencing of oat cultivar Ogle-C and transcriptome *de novo* assembly

The oat (*A. sativa* L.) genotype used for *de novo* transcriptome assembly was Ogle-C, derived from a single plant reselection of the cultivar ‘Ogle’ (Fox *et al.*, 2001). Developing dehulled seeds, collected at 7, 14, 21, and 28DAA (Gutierrez-Gonzalez, Wise, *et al.*, 2013), were the source of RNA. Two sets of libraries were constructed. First, libraries were constructed from RNA of all three biological replications from the four developmental stages (12 libraries, **Appendix S5**) and sequenced in paired-end mode with 100 cycles on the Illumina HiSeq 2000 machine as described previously (Gutierrez-Gonzalez, Tu, *et al.*, 2013). Second, longer reads were generated from a library constructed from a pool of RNA from the 4 developmental stages as described previously (Gutierrez-Gonzalez and Garvin, 2017), and sequenced on the Illumina MiSeq® platform using v3 chemistry generating 300 nt paired-end sequences (**Appendix S6**). Trimmomatic version 0.36 (Bolger *et al.*, 2014) was used to remove the first 12 nt, Illumina Truseq adaptor remnants and bases with an average quality within 4-bp sliding windows below a base quality value threshold of 20. A read was removed from the dataset if was shorter than 81 nt for HiSeq-generated sequenced and 181nt for Miseq sequences, respectively. Trimmed paired-end reads were assembled using Trinity v2.8.4 (Grabherr *et al.*, 2013) with default parameters.

### Validation of the *de novo* transcriptome assembly

We started with the longest isoform set from each Trinity “gene” (**Figure 1**). The longest isoform set was then aligned against the *Brachypodium distachyon* (UP000008810), *Hordeum vulgare* (UP000011116), and *Triticum aestivum* (UP000019116) predicted proteomes using NCBI blastx 2.7.1 (Camacho *et al.*, 2009) with an E-value cutoff of < 10^−10^. Trinity transcripts without any blast hits were aligned to the oat genome v1.0 (*Avena sativa* v1.0, http://avenagenome.org/, consisting of 63,455 scaffolds lacking annotation) using GMAP version 2018-07-04 (Wu and Watanabe, 2005) with parameter settings of >85% coverage and >85% identity, and all other parameters at default values. The unaligned Trinity transcript sets were searched against the UniRef100 database (Release: 2018_10, 07-Nov-2018) using NCBI blastx 2.7.1 with an E-value cutoff of < 10^−3^. For the transcripts that did not align to the draft oat genome, we extracted the best hit for each query sequence from the UniRef100 aligments and used taxonomic information to identify potential contaminant sequences. To assess the completeness of the oat transcriptome we evaluated the RTA using the BUSCO toolkit (Waterhouse *et al.*, 2018) using the Plantae lineage-specific single-copy orthologs (embryophyta odb 9) consisting of 1440 single copy orthologs.

### Experimental design, sample collection, 3’ RNA-Seq library construction, sequencing and metabolites chemical analysis of 22 oat lines

In 2016, we planted in the field and greenhouse 24 lines (**Table S2**) selected by clustering an oat diversity panel of 500 lines into 24 groups based on genotype and choosing the centroid of each cluster. This method of selection caused the lines to have low relatedness to each other, resulting in a genomic relationship matrix close to being diagonal (**Figure S12**).

In both trials, a randomized complete block design(RCBD) with two replicates was used (**Table S5**). Individual spikelets were tagged at anthesis and 10 spikelets were collected at 8, 13, 18, 23, 28 and 33 DAA. Primary florets were quickly dehulled on dry ice, then placed in liquid nitrogen and transferred to −80C freezer for storage. Two of the 24 lines without developing seeds collected at both sites were excluded. Of the 22 lines × 6 time points × 2 sites × 2 replicates = 528 possible samples, 419 samples with sufficient seed were randomly assigned to five 96-well plates for RNA extraction and 3’ RNA-Seq library construction using the same procedure as described by Kremling et al. (2018) at the Cornell University Sequencing facility. Pooled libraries were sequenced using Illumina NextSeq500 and HiSeq2000 with a 150 nt single-end run, v2 chemistry (**Appendix S7**).

After harvest, mature seeds were dehulled and analyzed with gas chromatography-mass spectrometry (GC-MS) and liquid chromatography–mass spectrometry (LC-MS) at the Proteomics and Metabolomics Facility at Colorado State University following Carlson *et al. (*2019).

### Quality trimming 3’ RNAseq reads, transcript quantification, and DGE analysis

BBMap version 37.50 (BBMap - Bushnell B. - sourceforge.net/projects/bbmap/) was used to remove adapter contamination, polyA sequences, and low quality sequences following a standard protocol described by Lexogen, Inc (QuantSeq User Guide) with slightly modified parameter settings of trimq=20, maq=20, and minlen=50 to retain reads with a minimum per base sequence quality score of 20 and minimum length of 50 nucleotides. After read quality control, expressed abundances were determined using Salmon version 0.12.0 (Patro *et al.*, 2017) and the RTA with default parameters. Samples with less than 0.5 million mapped reads and transcripts with less than two counts in at least ten samples were filtered out, leaving 59,815 transcripts for analysis. The filtered read count matrix was normalized by sequencing depth with a sample specific size factor implemented in DESseq2 version 1.22.2 (Love *et al.*, 2014). A PCA of samples was performed based on variance stabilized expression estimates using the vst function in DESeq2 package. The sample PCA plot showed two distinct clusters. We performed differential transcript expression analysis based on the major cluster of 326 samples (58,120 transcripts left after filtering those with less than two counts in at least ten samples) using the DESseq2 package. First, we performed a likelihood ratio test by comparing a full model (∼ genotype + time + genotype:time) against a reduced model (∼ genotype + time) to filter out transcripts showing a significant genotype-by-time interaction at FDR level of 0.1. This filter removed 424 transcripts, leaving 57,694 for subsequent analyses.

We performed a DGE analysis to identify transcripts differentially expressed between time points by controlling for the effect of different genotypes at FDR level of 0.05 using the standard method implemented in DESeq2 package. In order to understand how transcriptome expression correlated between time points, we averaged DESeq2 normalized read counts across samples within each time point for each transcript separately, and then applied a pairwise Spearman’s correlation analysis between time points. To identify global transcript expression patterns across time points common in all 22 oat lines, we constructed gene expression pattern sets consisting of DETs between any two adjacent time points. Based on the differential gene expression analysis results between any two adjacent time points, we partitioned all DETs in a single time interval into three categories including up-regulated, down-regulated and not differentially expressed, which were coded as “u”, “d”, and “0”, respectively. In this way, the expression pattern of each DET was coded as a string of five characters for the five time intervals. Finally, transcripts were classified into different temporal expression patterns based on their expression pattern codes.

### Heritability estimation of identified TCoE sets and simple linear regression between metabolites and GCoE sets

Variance components and heritability estimates of GCoE sets were based on the DESeq2 variance stabilized expression matrix with 397 samples and 59,815 transcripts after adjustment. We used the surrogate variable analysis (Leek *et al.*, 2012) to get an estimate of latent factors, and then the first latent factor was used to adjust for unwanted variation using the removeBatchEffect function implemented in R package limma (Ritchie *et al.*, 2015). For each transcript separately, the least square means (lsmean) of expression values of the 22 lines were estimated by the linear model ∼ Line + Location + Location/Replication + Time, generating an lsmean expression matrix. For each TCoE set, hierarchical clustering was used to partition transcripts into 4 to 20 sub-clusters based on the Euclidean distance of the lsmean expression matrix. The optimized number of subclusters of each TCoE set was determined by selecting the number of clusters that made heritabilities of all subclusters relatively high and with low variation. Using a TCoE set dependent and optimized number of sub-clusters is better than a uniform arbitrary number of sub-clusters applied to all TCoE sets because it allows different TCoE sets to have different genetic background partitions. For each GCoE set, PCA was applied to the lsmean expression matrix defined by the transcripts in that set, and scores of the first PC were extracted for the 22 oat lines. We fit nested models and performed a likelihood-ratio test: a full model, PC1score ∼ μ + Zu + e and a reduced model, PC1score ∼ μ + e. In the full model the random term u estimated the oat line additive effect with u ∼ N(0, K σ^2^_u_), where K was the genomic relationship matrix among the 22 oat lines (**Figure S12**) and σ^2^_u_ was the estimated additive genetic variance. For both models, the residual was distributed as e ∼ N(0, I σ^2^_e_), with I being an identity matrix. The heritability was estimated as σ^2^_u_ /(σ^2^_u_ + σ^2^_e_).

To test rigorously if the observed distribution of cluster sizes deviated from its expectation under the null distribution we used permutation testing. For a given TCoE set, expression of all genes were permuted relative to each other. The permuted matrix was then clustered to form eight clusters and the clusters ordered by size, but always dropping the smallest cluster. Permutation and clustering were repeated 1,000 times. The mean and covariance matrix among permuted cluster sizes were used to calculate the Mahalanobis distance of the non-permuted cluster size vector from the mean, and a corresponding p-value was calculated based on a Chi-Squared distribution with 7 degrees of freedom (Brereton, 2015; Wicklin 2012). This procedure was repeated for each of the 22 TCoE sets.

10 fatty acids, 282 and 529 metabolite features were obtained from targeted GC-MS, non-targeted GC-MS, non-targeted LC-MS analyses of mature seeds harvested from the two sites. A standard linear mixed model (∼Line + Location + Location/Replication + Location: Line) of the RCBD design was fitted for each metabolite using R package lme4 (Bates *et al.*, 2015), with all terms treated as random. The heritability was estimated as σ^2^_LINE_ /(σ^2^_LINE_ + σ^2^_LOCATION:LINE_/2 + σ^2^_e_/4). The metabolites with heritability > 0.4 were used as response variable in a simple linear regression with PC1 scores of each GCoE set as predictor. To compare the p-value obtained from real data against random sampling, for each metabolite and transcript abundance regression, we performed 100 permutations of PC1 scores of each GCoE set. Finally, p-values from permutation and non-permutation analyses were plotted.

### Transcriptome annotation and GO analysis

Functional annotation of the RTA was done following a standard workflow implemented in Trinotate v3.1.1 (Bryant *et al.*, 2017), which provided a comprehensive annotation including GO annotation assigned to each gene. To understand the biological functions behind the DETs between adjacent time points and those transcripts clustered to different temporal expression patterns, GO category over-representation analysis was performed using all transcripts of the RTA having at least one GO term as a background set with the R package of goseq v1.34.1 (Young *et al.*, 2010). Over-represented GO categories that were significant at FDR adjusted p-values of 0.01 were further plotted using the R package ComplexHeatmap 1.20.0 (Gu *et al.*, 2016).

## Acknowledgements

The authors thank Robin Buell for her valuable advice on *de novo* oat seed transcriptome assembly and comments on an earlier version of the manuscript; Daniel Ilut and Mandy Waters for their valuable advice on *de novo* transcriptome assembly; Jessica Schlueter for sharing unpublished oat genome sequences for validating the *de novo* assembled oat seed transcriptome; David Benscher and Amy Tamara Fox for help with planting field trials; Nicholas Kaczmar for assistance in collecting developing seeds; Sharon Mitchell, Jing Wu, Asha Jain for RNA extraction and library preparation. Funding for this research was provided by USDA-NIFA-AFRI grant number 2017-67007-26502 and by USDA-ARS project number 8062-21000-045-00D.

## Author contributions

J.J, M.A.G and M.E.S designed this project and supervised the research. H.H., J.J and M.A.G wrote the manuscript, and all co-authors were involved in editing the manuscript. H.H and X.L conducted the field experiment and collected samples for 3’ RNA sequencing. H.H. and J.J performed data analyses. J.J.G and D.F.G. generated full-length transcript RNA sequences of the oat cv. Ogle-C.

## Supporting information

Additional Supporting Information may be found online in the supporting information tab for this article:

**Figure S1** Transcript length distribution of the 131,457 transcripts included in the RTA.

**Figure S2** PCA plot of 397 samples with more than 0.5 million mapped reads based on the 500 transcripts with highest variance.

**Figure S3** Distribution of Pearson correlation coefficients of biological replicates from Greenhouse samples, Field samples and among samples across the two sites.

**Figure S4** Biological process (a), cellular compartments (b) and molecular function (c) GO terms enriched for differentially expressed transcript sets between adjacent time points. FDR adjusted p-values < 0.01 (in -log10 scale) were colored between blue and red, and cells without GO terms assigned were colored in gray.

**Figure S5** The 80 observed temporal transcript expression patterns identified from 25,971 differentially expressed transcripts between five pairs of adjacent time points.

**Figure S6** Correlation of transcript numbers (log scale) between each pair of symmetrical up- and down-regulated expression patterns. Each point represents a pair of symmetrical up- and down-regulated expression patterns. The number of transcripts in the up-regulated pattern on the x-axis and the number of transcript in the down-regulated pattern on the y-axis. Black points have one differential expression event, red points two, and green points three such events.

**Figure S7** GO categories enriched for 8 temporal transcript co-expression sets. FDR adjusted p-values < 0.01 (in -log10 scale) were colored between blue and red, and cells without GO terms assigned were colored in gray.

**Figure S8** Distribution of p-values of simple linear regression between 634 metabolites and PC1 scores of GCoE sets. Red boxes contained p-values from real data, and blue boxes contained p-values from 100 permutations.

**Figure S9** Transcript length distribution of the 9,817 transcripts that couldn’t be aligned to the UniRef100.

**Figure S10** clusters of 12 HiSeq samples based on expression profiles. Euclidean distances between samples were colored between dark blue and light blue.

**Figure S11** Pearson correlation coefficients of biological replicates from 12 HiSeq samples of cv.Ogle-C whose developing seeds were collected at 7, 14, 21, and 28 DAA.

**Figure S12** Heatmap of genomic relationship among 22 oat lines used in this study.

**Table S1** A comparison of BUSCOs plant gene completeness between the RTA in this study and the first version of *de novo* oat seed transcriptome assembly.

**Table S2** A list of 22 oat lines used in this study.

**Table S3** Chi-Square test for sub-cluster size distribution of the 22 temporal co-expression sets. **Table S4** A list of oat transcripts homologous to biosynthetic genes of avenanthremides and fatty acids from other oat cultivars and *Brachypodium distachyon.*

**Table S5** Detailed information of experimental design and 3’ RNASeq sample names.

**Appendix S1** A fasta file containing the 131,457 transcript sequences of the RTA.

**Appendix S2** An expression matrix of 59,815 transcripts by 397 samples. Sample names were coded as combination of Location, GID, DAA and block ID, which were described in Table S5. Expression abundances were normalized by sample specific size factor and then variance stabilization transformed using DESseq2.

**Appendix S3** Lists of differentially expressed genes between each pair of time points.

**Appendix S4** A file containing *de novo* assembled RTA transcripts annotation.

**Appendix S5** raw reads of the 12 libraries sequenced in paired-end mode with 100 cycles on the Illumina HiSeq 2000 platform.

**Appendix S6** raw reads of one library constructed from a pool of RNA from four developmental stages and sequenced by the Illumina MiSeq platform.

**Appendix S7** raw reads of 419 3’ RNASeq libraries sequenced by NextSeq500/HiSeq2000 with a 150 nt single-end run.

Datasets of appendices S1 to S7 are available on the CyVerse Data Commons. DOI: 10.25739/7y0n-de49 (Hu 2019). CyVerse Data Store file path: http://datacommons.cyverse.org/browse/iplant/home/shared/commons_repo/curated/HaixiaoHu_PBJOatTranscriptome_Oct2019.

## Footnotes

1 The term gene expression is used to indicate transcript abundance in this study.

## References

Banas, A., Debski, H., Banaś, W., Heneen, W.K., Dahlqvist, A., Bafor, M., et al. (2007) Lipids in grain tissues of oat (Avena sativa): Differences in content, time of deposition, and fatty acid composition. J. Exp. Bot., 58, 2463–2470.

Barthole, G., Lepiniec, L., Rogowsky, P.M., and Baud, S. (2012) Controlling lipid accumulation in cereal grains. Plant Sci., 185–186, 33–39.

Bates, D., Mächler, M., Bolker, B., and Walker, S. (2015) Fitting Linear Mixed-Effects Models Using lme4. J. Stat. Softw., 67.

Bian, J., Deng, P., Zhan, H., Wu, X., Nishantha, M.D.L.C., Yan, Z., et al. (2019) Transcriptional dynamics of grain development in barley (Hordeum vulgare L.). Int. J. Mol. Sci., 20, 1–16.

Bolger, A.M., Lohse, M., and Usadel, B. (2014) Trimmomatic: A flexible trimmer for Illumina sequence data. Bioinformatics, 30, 2114–2120.

Brereton, R.G. (2015) The chi squared and multinormal distributions. J. Chemom., 29, 9–12.

Bryant, D.M., Johnson, K., DiTommaso, T., Tickle, T., Couger, M.B., Payzin-Dogru, D., et al. (2017) A Tissue-Mapped Axolotl De Novo Transcriptome Enables Identification of Limb Regeneration Factors. Cell Rep., 18, 762–776.

Bryngelsson, S., Ishihara, A., and Dimberg, L.H. (2003) Levels of avenanthramides and activity of hydroxycinnamoyl-CoA:Hydroxyanthranilate N-hydroxycinnamoyl transferase (HHT) in steeped or germinated oat samples. Cereal Chem., 80, 356–360.

Camacho, C., Coulouris, G., Avagyan, V., Ma, N., Papadopoulos, J., Bealer, K., and Madden, T.L. (2009) BLAST+: architecture and applications. BMC Bioinformatics, 10, 421.

Carlson, M.O., Montilla-Bascon, G., Hoekenga, O.A., Tinker, N.A., Poland, J., Baseggio, M., et al. (2019) Multivariate Genome-wide Association Analyses Reveal the Genetic Basis of Seed Fatty Acid Composition in Oat (Avena sativa L.). bioRxiv, 589952.

Collins, F.W. (2011) Oat Phenolics: Biochemistry and Biological Functionality. In: Oats: Chemistry and Technology: Second Edition.

Fox, S.L., Jellen, E.N., Kianian, S.F., Rines, H.W., and Phillips, R.L. (2001) Assignment of RFLP linkage groups to chromosomes using monosomic F1 analysis in hexaploid oat. Theor. Appl. Genet., 102, 320–326.

Frey, K.J. and Holland, J.B. (1989) CROP BREEDING, GENETICS & CYTOLOGY Nine Cycles of Recurrent Selection for Increased Groat-Oil Content in Oat. 1636–1641.

Grabherr, M., Bj, H., Yassour, M., Levin, J., Thompson, D., Amit, I., et al. (2013) Trinity: reconstructing a full-length transcriptome without a genome from RNA-Seq data. Nat. Biotechnol., 29, 644–652.

Gu, Z., Eils, R., and Schlesner, M. (2016) Complex heatmaps reveal patterns and correlations in multidimensional genomic data. Bioinformatics, 32, 2847–2849.

Gutierrez-gonzalez, J.J. and Garvin, D.F. (2017) Oat. 1536, 209–221.

Gutierrez-Gonzalez, J.J., Tu, Z.J., and Garvin, D.F. (2013) Analysis and annotation of the hexaploid oat seed transcriptome. BMC Genomics, 14.

Gutierrez-Gonzalez, J.J., Wise, M.L., and Garvin, D.F. (2013) A developmental profile of tocol accumulation in oat seeds. J. Cereal Sci., 57, 79–83.

Hirsch, C.N., Foerster, J.M., Johnson, J.M., Sekhon, R.S., Muttoni, G., Vaillancourt, B., et al. (2014) Insights into the Maize Pan-Genome and Pan-Transcriptome. Plant Cell, 26, 121–135.

Hoffman, L.A. (2011) World production and use of oats. Oat Crop, 34–61.

Haixiao Hu (2019) Heritable temporal gene expression patterns correlate with metabolomic seed content in developing hexaploid oat seed - Supporting information. CyVerse Data Commons. DOI 10.25739/7y0n-de49

Ilut, D.C., Sanchez, P.L., Costich, D.E., Friebe, B., Coffelt, T.A., Dyer, J.M., et al. (2015) Genomic diversity and phylogenetic relationships in the genus Parthenium (Asteraceae). Ind. Crops Prod., 76, 920–929.

Krafft, C., Dietzek, B., and Popp, J. (2011) Biomedical Imaging Based on Vibrational Spectroscopy. Opt. Digit. Image Process. Fundam. Appl., 717–737.

Kremling, K.A.G., Chen, S.Y., Su, M.H., Lepak, N.K., Romay, M.C., Swarts, K.L., et al. (2018) Dysregulation of expression correlates with rare-allele burden and fitness loss in maize. Nature, 555, 520–523.

Leek, J.T., Johnson, W.E., Parker, H.S., Jaffe, A.E., and Storey, J.D. (2012) The SVA package for removing batch effects and other unwanted variation in high-throughput experiments. Bioinformatics, 28, 882–883.

Li, G., Wang, D., Yang, R., Logan, K., Chen, H., Zhang, S., et al. (2014) Temporal patterns of gene expression in developing maize endosperm identified through transcriptome sequencing. Proc. Natl. Acad. Sci., 111, 7582–7587.

Li, Y., Fu, X., Zhao, M., Zhang, W., Li, B., An, D., et al. (2018) A Genome-wide View of Transcriptome Dynamics During Early Spike Development in Bread Wheat. Sci. Rep., 8, 1–16.

Liu, K.S. (2011) Comparison of Lipid Content and Fatty Acid Composition and Their Distribution within Seeds of 5 Small Grain Species. J. Food Sci., 76, 334–342.

Love, M.I., Anders, S., Kim, V., and Huber, W. (2016) RNA-Seq workflow: gene-level exploratory analysis and differential expression. F1000Research, 4, 1070.

Love, M.I., Huber, W., and Anders, S. (2014) Moderated estimation of fold change and dispersion for RNA-seq data with DESeq2. Genome Biol., 15, 550.

Matsukawa, T., Isobe, T., Ishihara, A., and Iwamura, H. (2000) Occurrence of avenanthramides and hydroxycinnamoyl-CoA:hydroxyanthranilate N-hydroxycinnamoyltransferase activity in oat seeds. Zeitschrift fur Naturforsch. - Sect. C J. Biosci., 55, 30–36.

Moll, P., Ante, M., Seitz, A., and Reda, T. (2014) QuantSeq 3′ mRNA sequencing for RNA quantification. Nat. Methods, 11, i–iii.

Patro, R., Duggal, G., Love, M.I., Irizarry, R.A., and Kingsford, C. (2017) Salmon provides fast and bias-aware quantification of transcript expression. Nat. Methods, 14, 417–419.

Peterson, D.M. and Dimberg, L.H. (2008) Avenanthramide concentrations and hydroxycinnamoyl-CoA:hydroxyanthranilate N-hydroxycinnamoyltransferase activities in developing oats. J. Cereal Sci., 47, 101–108.

Rangan, P., Furtado, A., and Henry, R.J. (2017) The transcriptome of the developing grain: A resource for understanding seed development and the molecular control of the functional and nutritional properties of wheat. BMC Genomics, 18, 1–9.

Rasane, P., Jha, A., Sabikhi, L., Kumar, A., and Unnikrishnan, V.S. (2013) Nutritional advantages of oats and opportunities for its processing as value added foods - a review. J. Food Sci. Technol., 52, 662–675.

Ritchie, M.E., Phipson, B., Wu, D., Hu, Y., Law, C.W., Shi, W., and Smyth, G.K. (2015) limma powers differential expression analyses for RNA-sequencing and microarray studies. Nucleic Acids Res., 43, e47.

Ruuska, S.A. (2002) Contrapuntal Networks of Gene Expression during Arabidopsis Seed Filling. Plant Cell Online, 14, 1191–1206.

Simão, F.A., Waterhouse, R.M., Ioannidis, P., Kriventseva, E. V., and Zdobnov, E.M. (2015) BUSCO: Assessing genome assembly and annotation completeness with single-copy orthologs. Bioinformatics, 31, 3210–3212.

Tripathi, V., Singh, A., and Ashraf, M.T. (2018) Avenanthramides of oats: Medicinal importance and future perspectives. Pharmacogn. Rev.

Tzfadia, O., Bocobza, S., Defoort, J., Almekias-Siegl, E., Panda, S., Levy, M., et al. (2018) The ‘TranSeq’ 3′-end sequencing method for high-throughput transcriptomics and gene space refinement in plant genomes. Plant J., 96, 223–232.

USDA (2019) Grain: World Markets and Trade Competitive Pricing Suggests Rebound in EU Wheat Exports.

Wan, Y., Poole, R.L., Huttly, A.K., Toscano-Underwood, C., Feeney, K., Welham, S., et al. (2008) Transcriptome analysis of grain development in hexaploid wheat. BMC Genomics, 9, 1–16.

Wang, Y., Lysøe, E., Armarego-Marriott, T., Erban, A., Paruch, L., Van Eerde, A., et al. (2018) Transcriptome and metabolome analyses provide insights into root and root-released organic anion responses to phosphorus deficiency in oat. J. Exp. Bot., 69, 3759–3771.

Waterhouse, R.M., Seppey, M., Simao, F.A., Manni, M., Ioannidis, P., Klioutchnikov, G., et al. (2018) BUSCO applications from quality assessments to gene prediction and phylogenomics. Mol. Biol. Evol., 35, 543–548.

Wise, M. (2017) Tissue Distribution of Avenanthramides and Gene Expression of Hydroxycinnamoyl-CoA:hydroxyanthranilate N-hydroxycinnamoyl Transferase (HHT) in Benzothiadiazole Treated. Can. J. Plant Sci., 456, 444–456.

Wu, B., Hu, Y., Huo, P., Zhang, Q., Chen, X., and Zhang, Z. (2017) Transcriptome analysis of hexaploid hulless Oat in response to salinity stress. PLoS One, 12, 1–16.

Wu, T.D. and Watanabe, C.K. (2005) GMAP: A genomic mapping and alignment program for mRNA and EST sequences. Bioinformatics, 21, 1859–1875.

Yang, Q., Xuan Trinh, H., Imai, S., Ishihara, A., Zhang, L., Nakayashiki, H., et al. (2004) Analysis of the Involvement of Hydroxyanthranilate Hydroxycinnamoyltransferase and Caffeoyl-CoA 3-O -Methyltransferase in Phytoalexin Biosynthesis in Oat. Mol. Plant-Microbe Interact., 17, 81–89.

Yi, F., Gu, W., Chen, J., Song, N., Gao, X., Zhang, X., et al. (2019) High Temporal-Resolution Transcriptome Landscape of Early Maize Seed Development. Plant Cell, 31, 974–992.

Young, M.D., Wakefield, M.J., Smyth, G.K., and Oshlack, A. (2010) Gene ontology analysis for RNA-seq: accounting for selection bias GOseq GOseq is a method for GO analysis of RNA-seq data that takes into account the length bias inherent in RNA-seq. Genome Biol., 11.

Zhang, R., Tucker, M.R., Burton, R.A., Shirley, N.J., Little, A., Morris, J., et al. (2016) The dynamics of transcript abundance during cellularisation of developing barley endosperm. Plant Physiol., 170, 1549–1565.

