## Supporting Information for "Heritable temporal gene expression patterns correlate with metabolomic seed content in developing hexaploid oat seed"

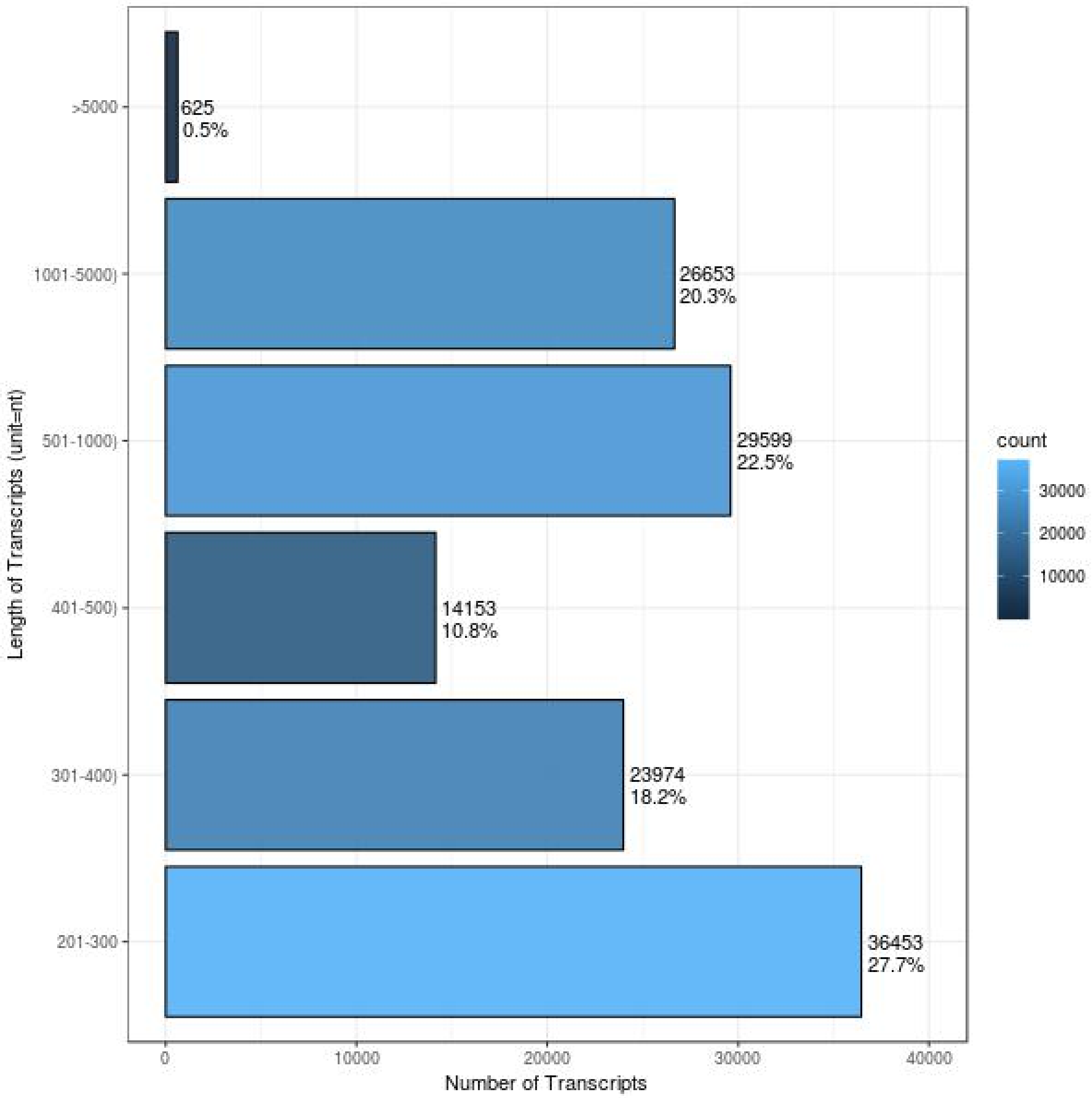


**Figure S1** Transcript length distribution of the 131,457 transcripts included in the RTA

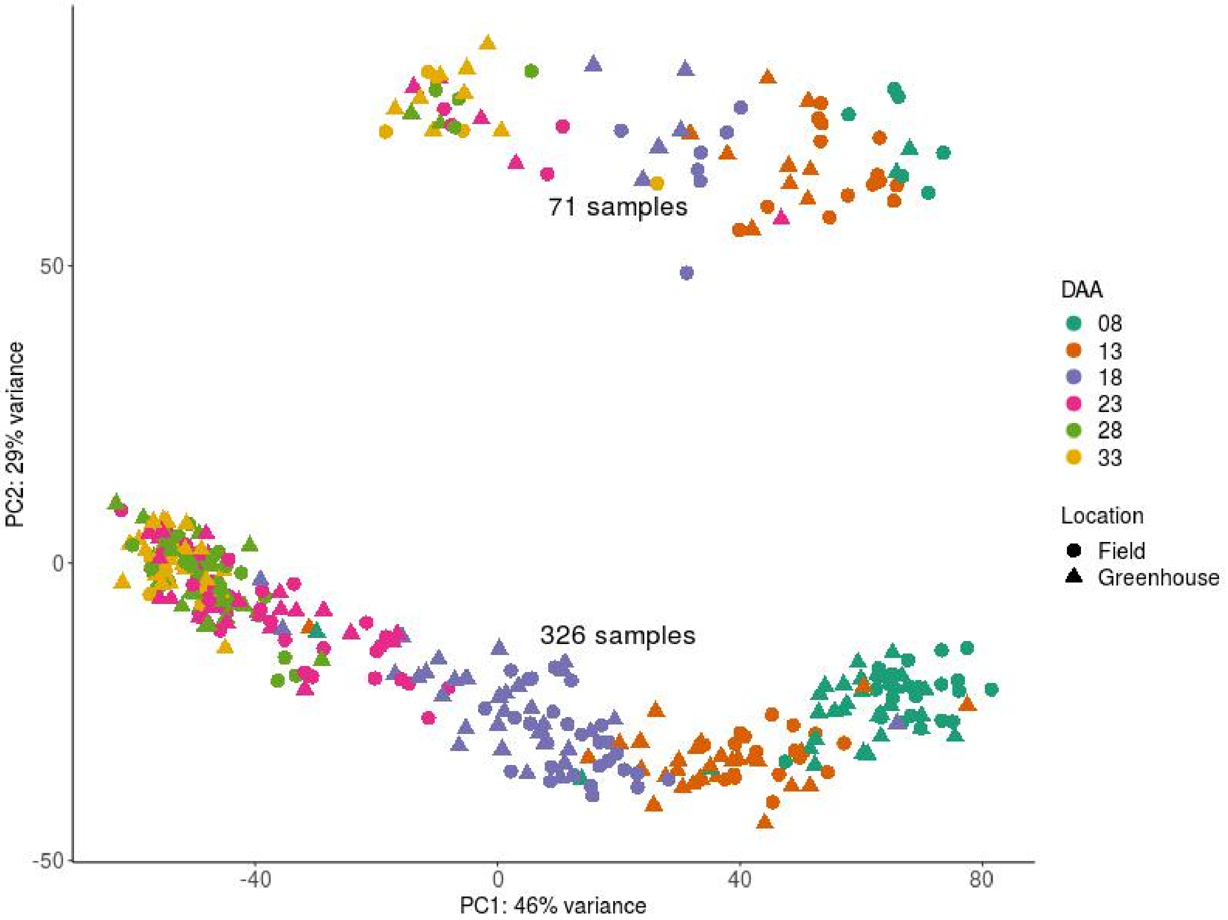


**Figure S2** PCA plot of 397 samples with more than 0.5 million mapped reads based on the 500 transcripts with highest variance.

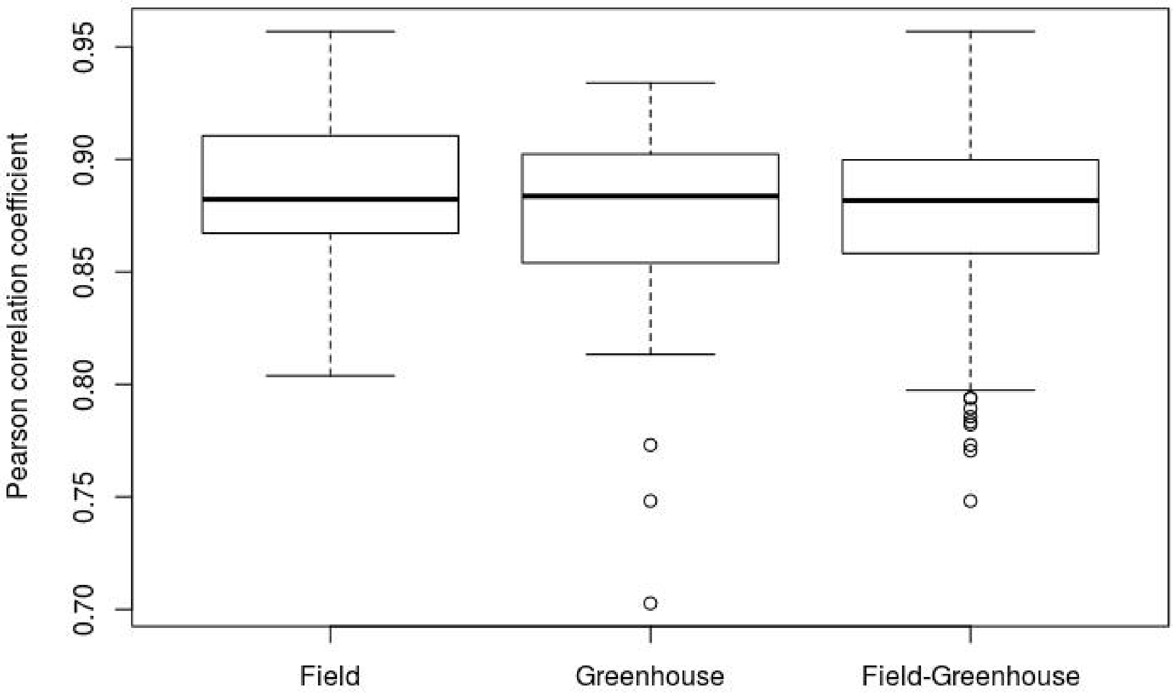


**Figure S3** Distribution of Pearson correlation coefficients of biological replicates from Greenhouse samples, Field samples and among samples across the two sites

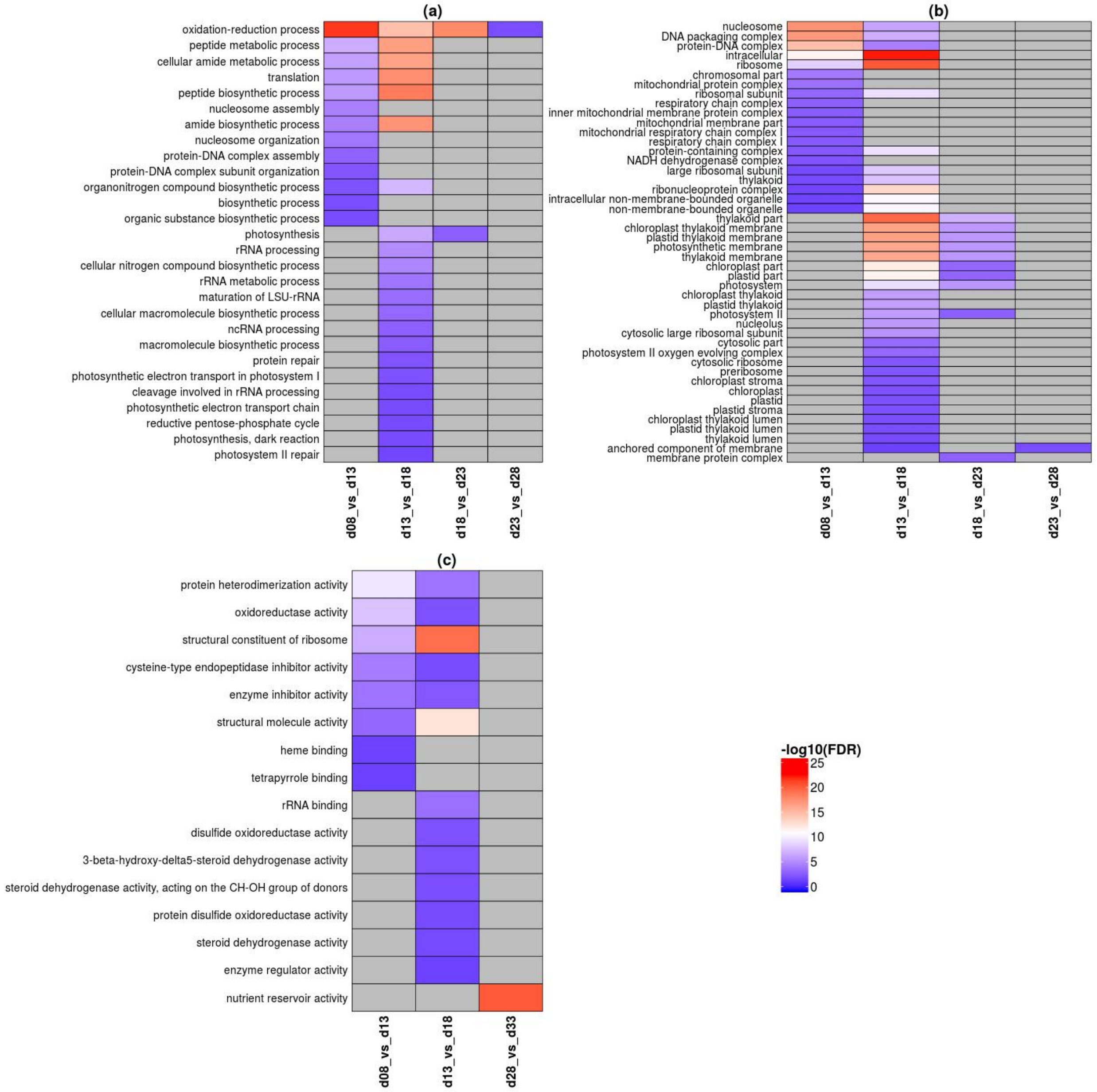


**Figure S4** Biological process (a), cellular compartments (b) and molecular function (c) GO terms enriched for differentially expressed transcript sets between adjacent time points. FDR adjusted p-values < 0.01 (in -log10 scale) were colored between blue and red, and cells without GO terms assigned were colored in gray.

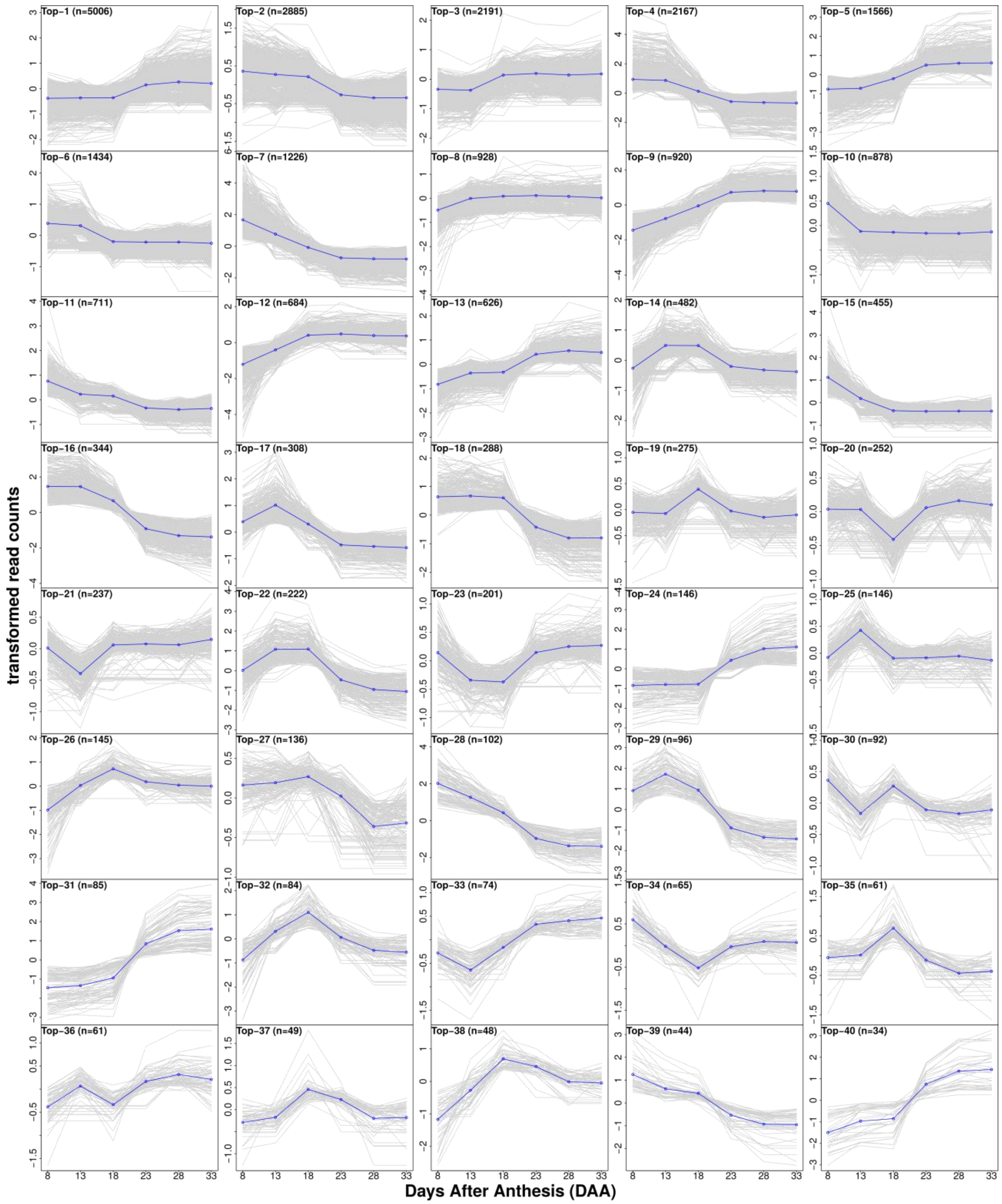


**Figure S5** Top 1-40 of the 80 observed temporal transcript expression patterns identified from 25,971 differentially expressed transcripts between five pairs of adjacent time points

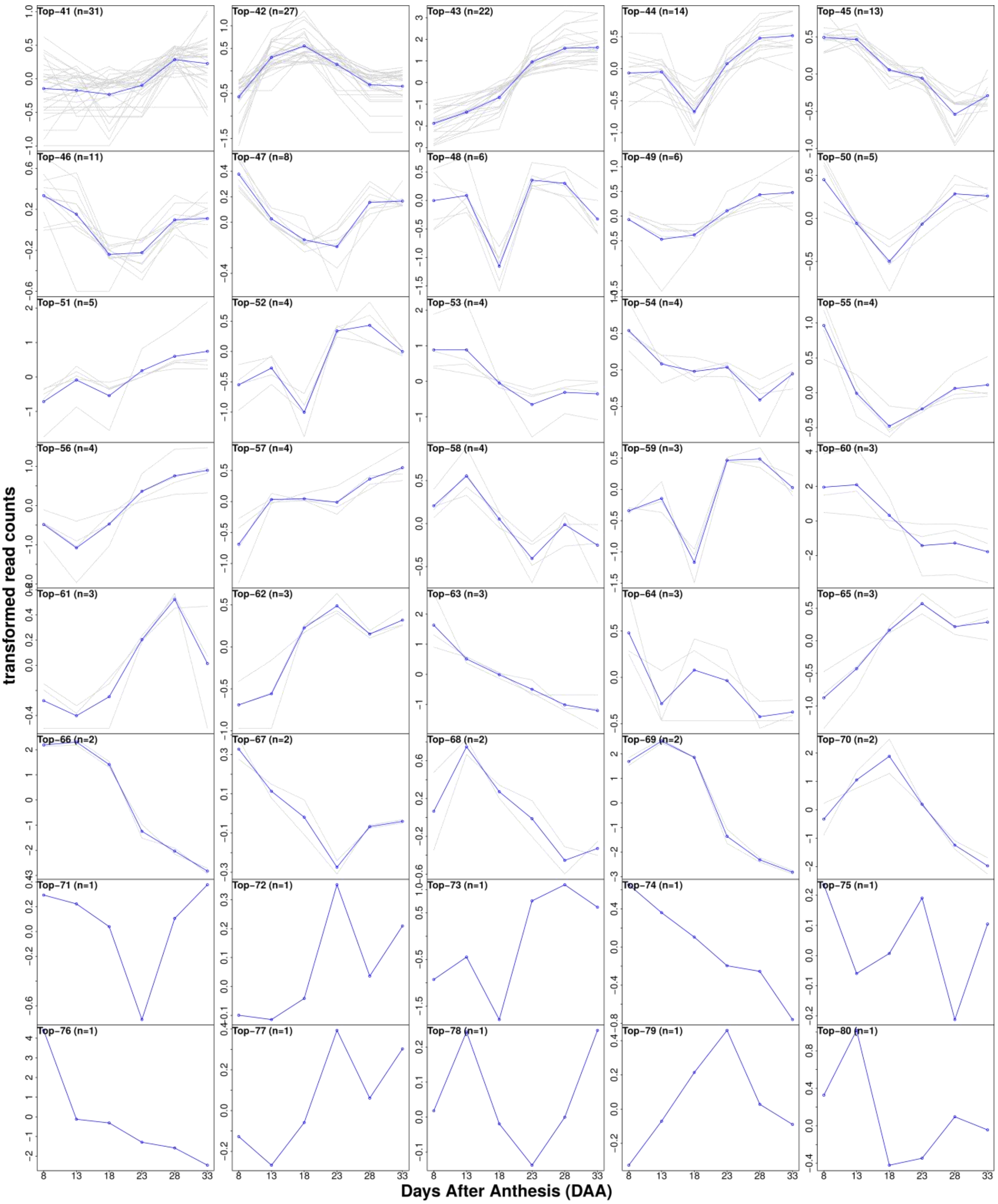


**Figure S5 (continued)** Top 41-80 of the 80 observed temporal transcript expression patterns identified from 25,971 differentially expressed transcripts between five pairs of adjacent time points

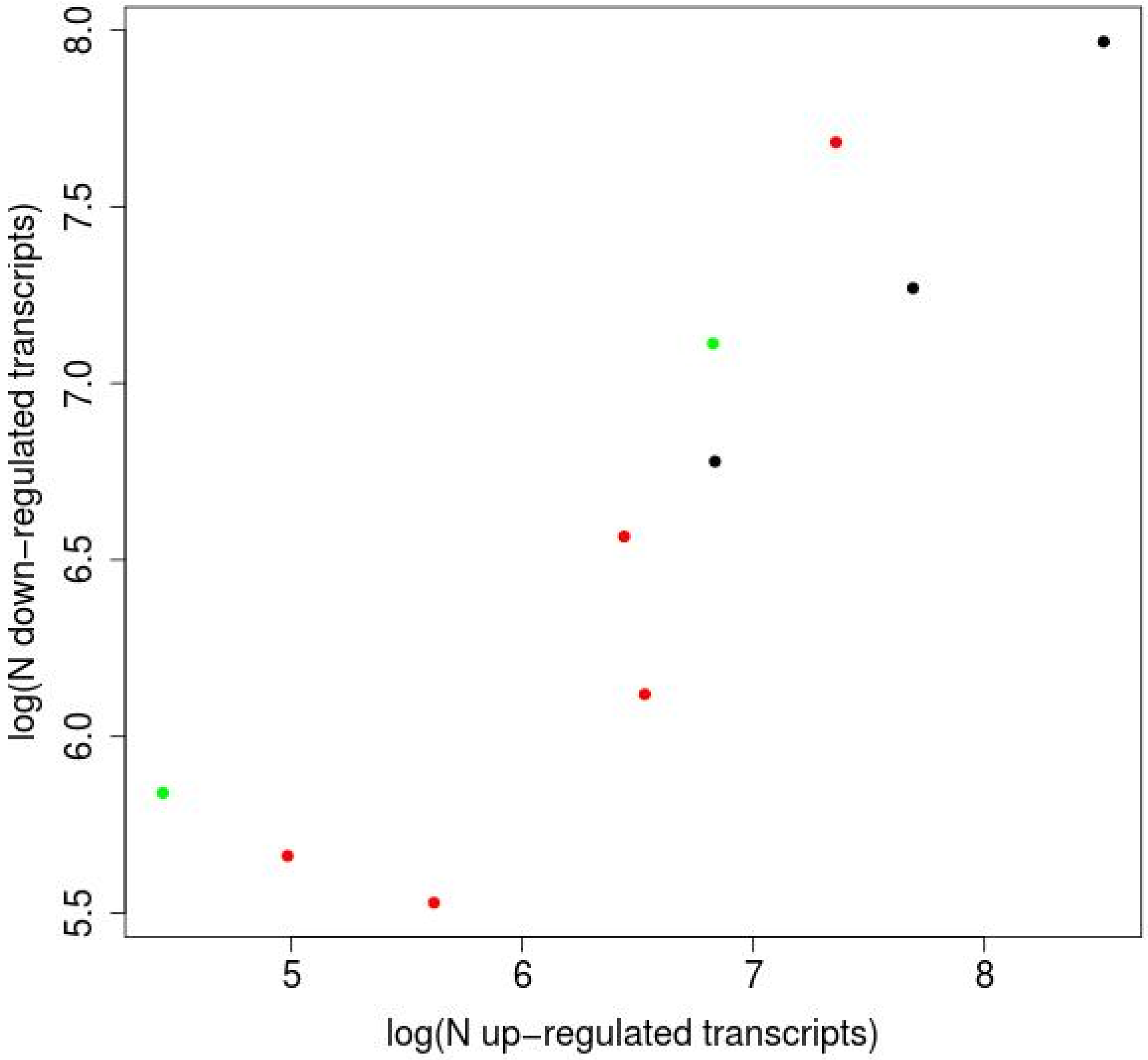


**Figure S6** Correlation of transcript numbers (log scale) between each pair of symmetrical up- and down-regulated expression patterns. Each point represents a pair of symmetrical up- and down-regulated expression patterns. The number of transcripts in the up-regulated pattern on the x-axis and the number of transcript in the down-regulated pattern on the y-axis. Black points have one differential expression event, red points two, and green points three such events.

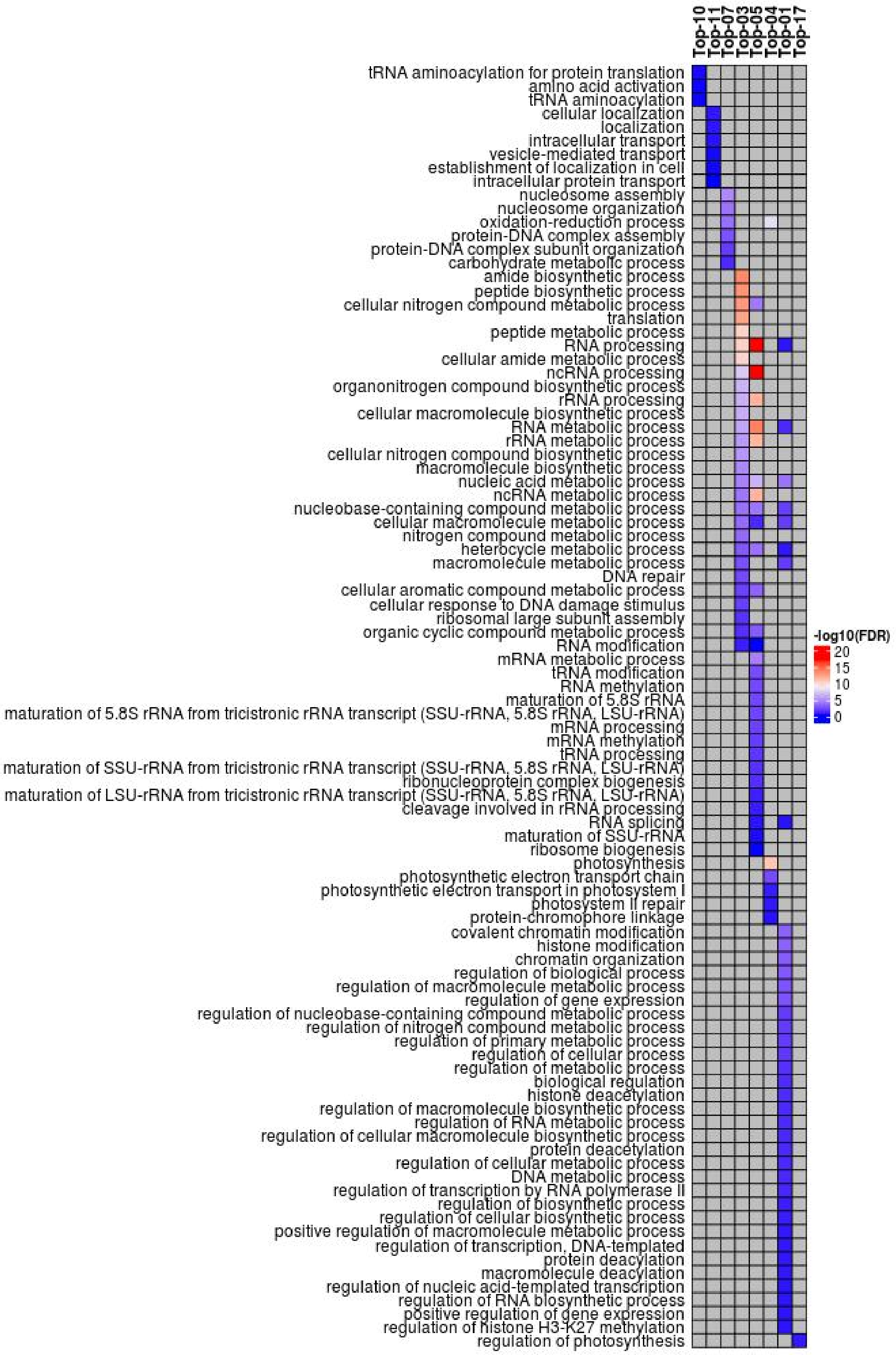


**Figure S7** GO categories enriched for 8 temporal transcript co-expression sets. FDR adjusted p-values < 0.01 (in -log10 scale) were colored between blue and red, and cells without GO terms assigned were colored in gray.

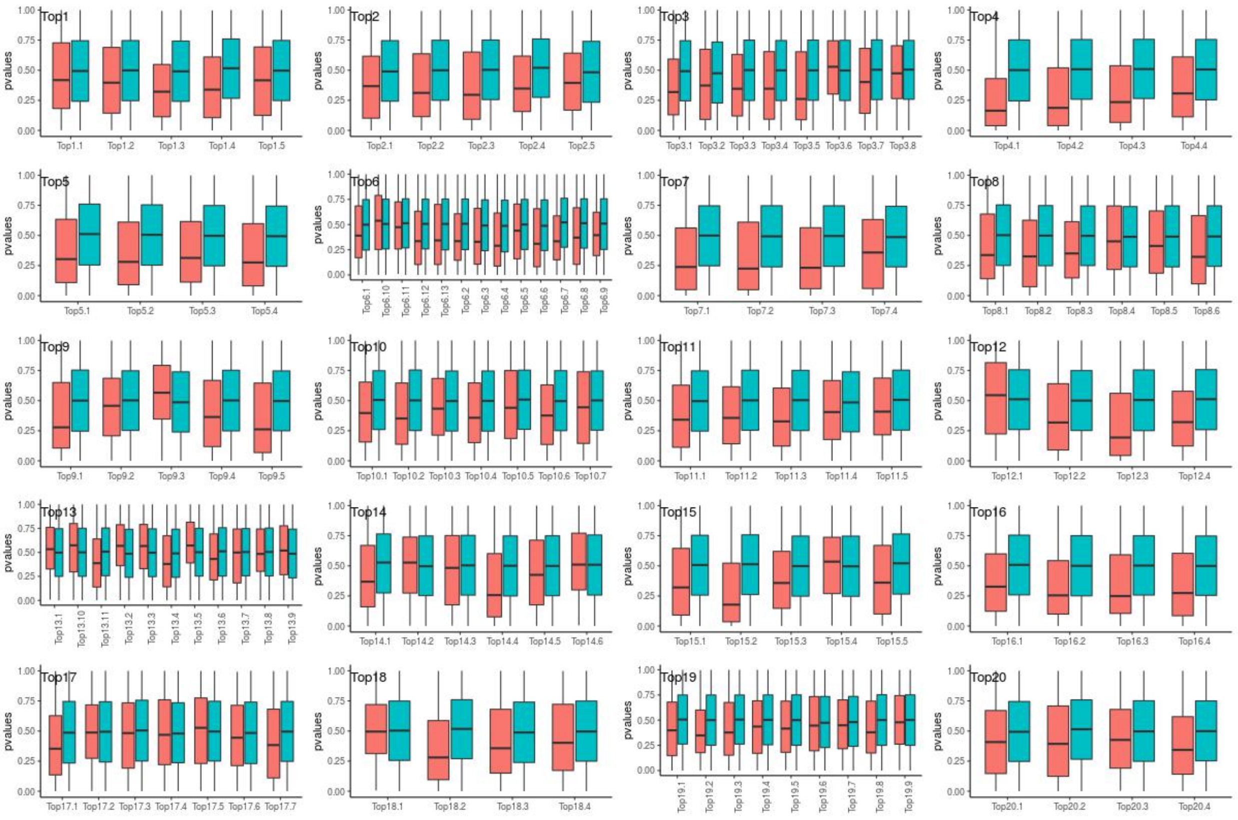

**Figure S8** Distribution of p-values of simple linear regression between 634 metabolites and PC1 scores of GCoE sets. Red boxes contained p-values from real data, and blue boxes contained p-values from 100 permutations.

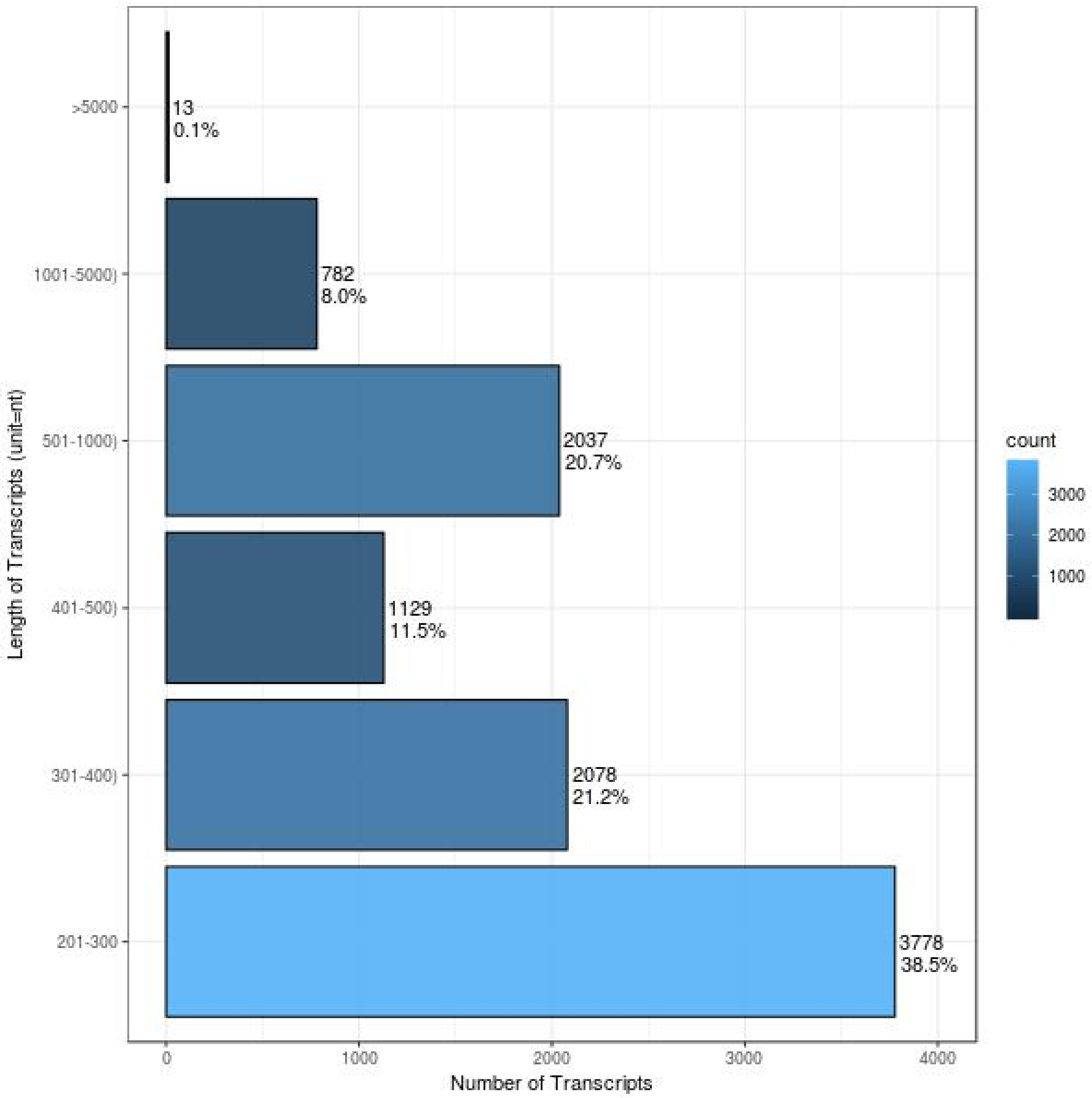


**Figure S9** Transcript length distribution of the 9,817 transcripts that couldn't be aligned to the UniRef100

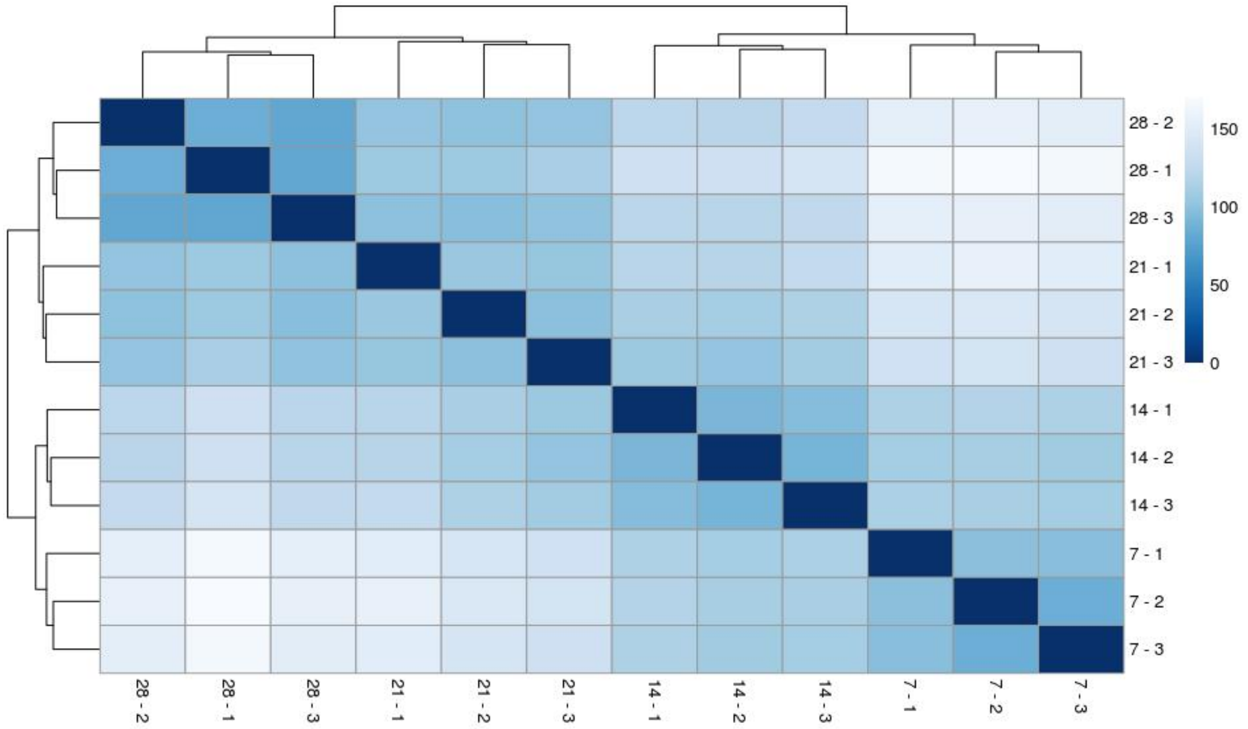


**Figure S10** clusters of 12 HiSeq samples based on expression profiles. Euclidean distances between samples were colored between dark blue and light blue.

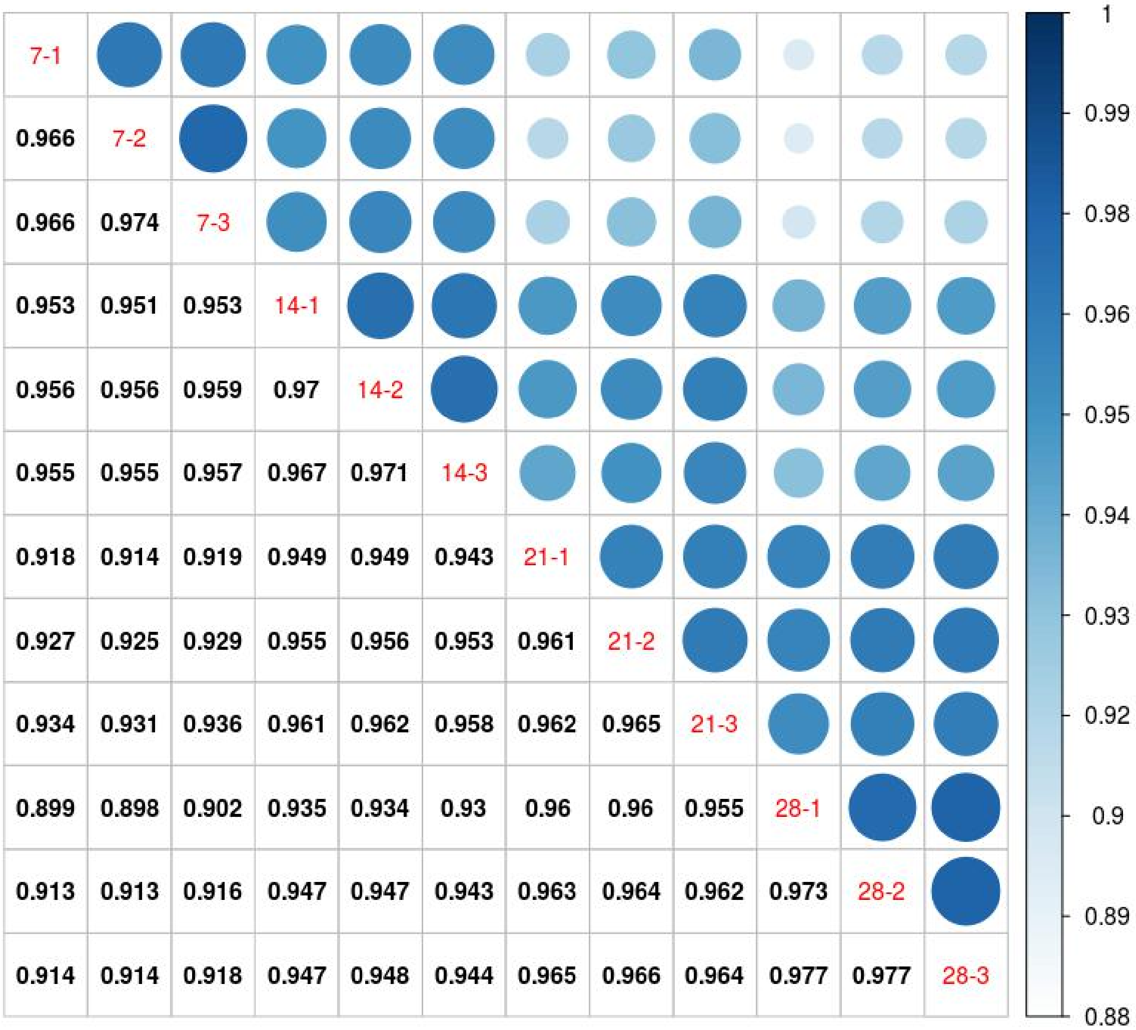


**Figure S11** Pearson correlation coefficients of biological replicates from 12 HiSeq samples of cv.Ogle-C whose developing seeds were collected at 7, 14, 21, and 28 DAA

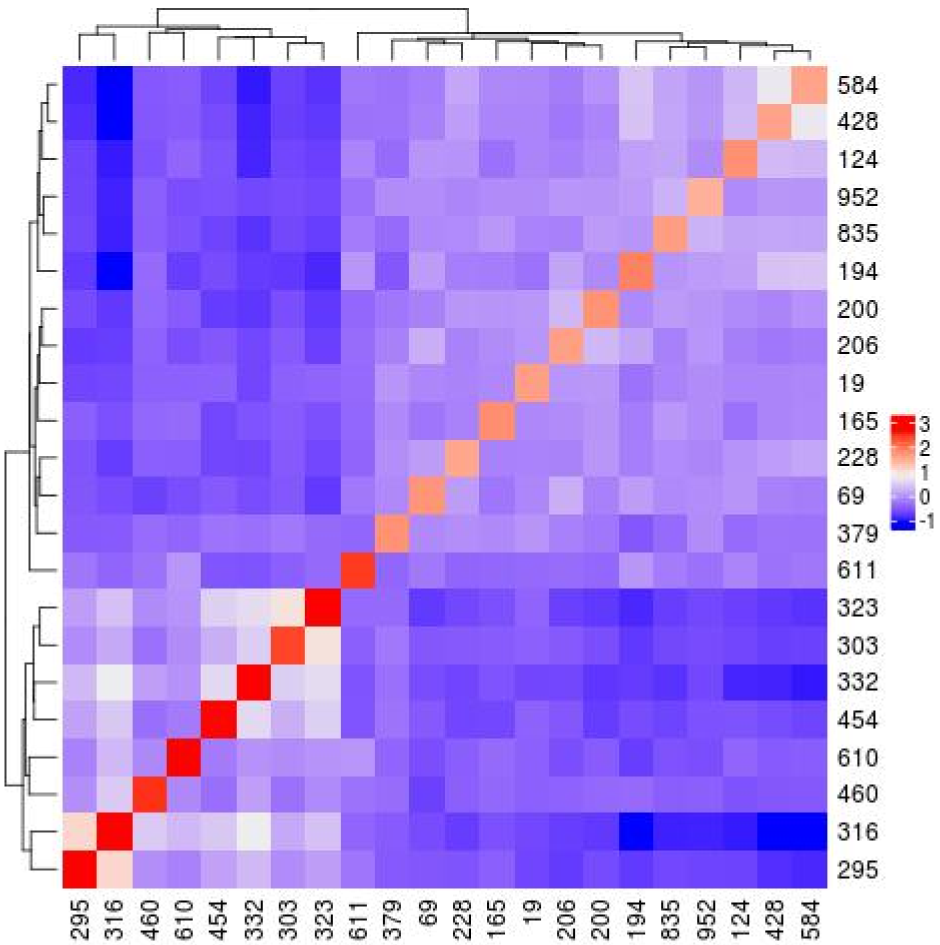


**Figure S12** Heatmap of genomic relationship among 22 oat lines used in this study

**Table S1** A comparison of BUSCOs plant gene completeness between the RTA in this study and the first version of *de novo* oat seed transcriptome assembly (*dn*OST, Gutierrez-Gonzalez et al. 2013)

| BUSCO Statistics | the RTA | *dn*OST |
|---|---|---|
| Complete BUSCOs | 1212 (84.17%) | 412 (28.61%) |
| Complete and single-copy BUSCOs | 1188 (82.50%) | 116 (8.06%) |
| Complete and duplicated BUSCOs | 24 (1.67%) | 296 (20.56%) |
| Fragmented BUSCOs | 148 (10.28%) | 201 (13.96%) |
| Missing BUSCOs | 80 (5.56%) | 827 (57.43%) |
| Total BUSCO groups searched | 1440 | 1440 |

**Table S2** A list of 22 oat lines used in this study

| GID | T3Oat_Name[1] |
|---|---|
| 19 | IL05-8515 |
| 69 | X8826-1 |
| 124 | SHERWOOD |
| 165 | MN08130 |
| 194 | ANDREW |
| 200 | EXCEL |
| 206 | WOODBURN |
| 228 | 95AB12770 |
| 295 | TIFT |
| 303 | FL0238BSB-22 |
| 316 | TX02U7479 |
| 323 | TX07CS1402 |
| 332 | HORIZON270 |
| 428 | PI344841 |
| 454 | PERDEBERG |
| 460 | CIAV5389 |
| 584 | ONOHOJSKIJ_A-547 |
| 610 | QUALITY\|PI289587 |
| 611 | CIAV6218 |
| 835 | IA111003 |
| 952 | M2-2 |
| 379 | CORRAL |

[1]T3Oat_Name= line names were consistent with that on T3/Oat (https://triticeaetoolbox.org/oat/)

**Table S3** Chi-Square test for sub-cluster size distribution of the 22 temporal co-expression sets. At significant level of 0.05, the critical p-value after Bonferroni correction is 0.05/22=0.00227.

| Temporal co-expression set* | P-value |
|---|---|
| Top-8 | 0.994548451 |
| Top-10 | 0.999286221 |
| Top-12 | 0.497109215 |
| Top-15 | 0.003939038 |
| Top-13 | 0.108831006 |
| Top-11 | 0.344207141 |
| Top-9 | 0.380203442 |
| Top-7 | 0.608797221 |
| Top-3 | 0.895528366 |
| Top-6 | 0.419936316 |
| Top-5 | 0.273601551 |
| Top-4 | 0.594794097 |
| Top-31 | 0.021221027 |
| Top-16 | 0.133281338 |
| Top-1 | 0.637929437 |
| Top-2 | 0.83840654 |
| Top-24 | 0.744979403 |
| Top-18 | 0.210760149 |
| Top-19 | 0.980442768 |
| Top-20 | 0.35296887 |
| Top-14 | 0.962213032 |
| Top-17 | 0.549011073 |

*The temporal co-expression sets were ordered the same as in Figure 4.

**Table S4** A list of oat transcripts homologous to biosynthetic genes of avenanthremides and fatty acids from other oat cultivars and *Brachypodium distachyon*

| Genename | Source | Transcript name | reference | query_len[a] | ref_len[b] | aln_length[c] | pct_ident[d] |
|---|---|---|---|---|---|---|---|
| **Avenanthramides biosynthetic genes** | | | | | | | |
| CCoA3H | Arabidopsis | TRINITY_DN6473_c0_g1_i1 | NM_180006.2 | 1942 | 1885 | 1362 | 67.621 |
| CCoA3H | Brachypodium | TRINITY_DN6473_c0_g1_i1 | KQK05634 | 1942 | 2031 | 1954 | 86.131 |
| CCoAOMT | Oat | TRINITY_DN3508_c0_g1_i7 | AB076979.1 | 1238 | 622 | 604 | 99.834 |
| HHT1 | Oat | TRINITY_DN5172_c0_g1_i4 | AB076980.1 | 1598 | 1755 | 1343 | 79.449 |
| HHT2 | Oat | TRINITY_DN5172_c0_g1_i4 | AB076981.1 | 1598 | 1770 | 1343 | 78.555 |
| HHT3 | Oat | TRINITY_DN5172_c0_g1_i4 | AB076982.1 | 1598 | 1659 | 1343 | 79.598 |
| HHT4 | Oat | TRINITY_DN5172_c0_g1_i4 | AB076983.1 | 1598 | 1100 | 1088 | 97.151 |
| **Fatty Acids biosynthetic genes** | | | | | | | |
| ACCase | Brachypodium | TRINITY_DN146_c0_g1_i8 | BRADI_5g03860 | 7812 | 8783 | 7598 | 88.694 |
| DGAT1/TAG1 | Brachypodium | TRINITY_DN151_c0_g1_i4 | BRADI_1g37750 | 2878 | 3009 | 925 | 72.757 |
| FAB1/KAS2 | Brachypodium | TRINITY_DN207_c0_g1_i1 | BRADI_1g60300 | 2876 | 2498 | 1851 | 87.088 |
| FAB2 | Brachypodium | TRINITY_DN2740_c0_g1_i5 | BRADI_2g58930 | 5078 | 1792 | 1610 | 86.957 |
| FAD2 | Brachypodium | TRINITY_DN268_c0_g1_i4 | BRADI_3g53370 | 1883 | 2351 | 1665 | 85.526 |
| FAD3 | Brachypodium | TRINITY_DN10818_c0_g1_i3 | BRADI_1g65580 | 1198 | 2817 | 1195 | 82.343 |
| FAE1/KCS18 | Brachypodium | TRINITY_DN1011_c0_g1_i5 | BRADI_2g16050 | 1990 | 2481 | 1447 | 81.064 |
| FATB | Brachypodium | TRINITY_DN2629_c0_g1_i1 | BRADI_1g51170 | 2394 | 4407 | 1368 | 88.085 |
| GPAT9 | Brachypodium | TRINITY_DN5048_c0_g1_i7 | BRADI_1g25790 | 1490 | 1540 | 1460 | 86.37 |
| LPCAT1 | Brachypodium | TRINITY_DN3352_c0_g1_i1 | BRADI_3g51577 | 1846 | 1671 | 1634 | 88.433 |
| PAH1 | Brachypodium | TRINITY_DN1042_c0_g1_i6 | BRADI_2g23040 | 4252 | 4162 | 2485 | 82.817 |
| PDAT1 | Brachypodium | TRINITY_DN2921_c0_g1_i1 | BRADI_4g31540 | 2494 | 2601 | 2531 | 86.448 |
| WRI1 | Brachypodium | TRINITY_DN331_c0_g1_i1 | BRADI_4g43877 | 3900 | 2035 | 1223 | 80.376 |

[a]query_len= query sequence length;

[b]ref_len = reference sequence length;

[c]aln_length= alignment length;

[d]pct_ident= percent identity

**Table S5** Detailed information of experimental design and 3' RNASeq sample names

| Location | Block | Plot | GID | T3Oat_Name[1] | RNASeq_Samplename[2] |
|---|---|---|---|---|---|
| Greenhouse | 1 | 1 | 206 | WOODBURN | G001 |
| Greenhouse | 1 | 2 | 69 | X8826-1 | G002 |
| Greenhouse | 1 | 3 | 124 | SHERWOOD | G003 |
| Greenhouse | 1 | 4 | 316 | TX02U7479 | G004 |
| Greenhouse | 1 | 5 | 19 | IL05-8515 | G005 |
| Greenhouse | 1 | 7 | 332 | HORIZON270 | G007 |
| Greenhouse | 1 | 8 | 835 | IA111003 | G008 |
| Greenhouse | 1 | 9 | 428 | PI344841 | G009 |
| Greenhouse | 1 | 10 | 379 | CORRAL | G010 |
| Greenhouse | 1 | 11 | 165 | MN08130 | G011 |
| Greenhouse | 1 | 13 | 228 | 95AB12770 | G013 |
| Greenhouse | 1 | 14 | 610 | QUALITY\|PI289587 | G014 |
| Greenhouse | 1 | 15 | 952 | M2-2 | G015 |
| Greenhouse | 1 | 16 | 460 | CIAV5389 | G016 |
| Greenhouse | 1 | 17 | 323 | TX07CS1402 | G017 |
| Greenhouse | 1 | 18 | 454 | PERDEBERG | G018 |
| Greenhouse | 1 | 19 | 200 | EXCEL | G019 |
| Greenhouse | 1 | 20 | 303 | FL0238BSB-22 | G020 |
| Greenhouse | 1 | 21 | 584 | ONOHOJSKIJ_A-547 | G021 |
| Greenhouse | 1 | 22 | 194 | ANDREW | G022 |
| Greenhouse | 1 | 24 | 611 | CIAV6218 | G024 |
| Greenhouse | 1 | 25 | 295 | TIFT | G025 |
| Greenhouse | 2 | 26 | 19 | IL05-8515 | G026 |
| Greenhouse | 2 | 28 | 124 | SHERWOOD | G028 |
| Greenhouse | 2 | 29 | 332 | HORIZON270 | G029 |
| Greenhouse | 2 | 30 | 206 | WOODBURN | G030 |
| Greenhouse | 2 | 31 | 584 | ONOHOJSKIJ_A-547 | G031 |
| Greenhouse | 2 | 32 | 303 | FL0238BSB-22 | G032 |
| Greenhouse | 2 | 33 | 428 | PI344841 | G033 |
| Greenhouse | 2 | 34 | 611 | CIAV6218 | G034 |
| Greenhouse | 2 | 35 | 460 | CIAV5389 | G035 |
| Greenhouse | 2 | 36 | 228 | 95AB12770 | G036 |
| Greenhouse | 2 | 37 | 454 | PERDEBERG | G037 |
| Greenhouse | 2 | 38 | 69 | X8826-1 | G038 |
| Greenhouse | 2 | 39 | 835 | IA111003 | G039 |
| Greenhouse | 2 | 40 | 610 | QUALITY\|PI289587 | G040 |
| Greenhouse | 2 | 41 | 323 | TX07CS1402 | G041 |

| Location | Block | Plot | GID | T3Oat_Name[1] | RNASeq_Samplename[2] |
|---|---|---|---|---|---|
| Greenhouse | 2 | 42 | 952 | M2-2 | G042 |
| Greenhouse | 2 | 43 | 165 | MN08130 | G043 |
| Greenhouse | 2 | 45 | 295 | TIFT | G045 |
| Greenhouse | 2 | 47 | 194 | ANDREW | G047 |
| Greenhouse | 2 | 48 | 379 | CORRAL | G048 |
| Greenhouse | 2 | 49 | 316 | TX02U7479 | G049 |
| Greenhouse | 2 | 50 | 200 | EXCEL | G050 |
| Field | 1 | 803 | 124 | SHERWOOD | C803 |
| Field | 1 | 804 | 200 | EXCEL | C804 |
| Field | 1 | 806 | 303 | FL0238BSB-22 | C806 |
| Field | 1 | 807 | 316 | TX02U7479 | C807 |
| Field | 1 | 808 | 428 | PI344841 | C808 |
| Field | 1 | 809 | 206 | WOODBURN | C809 |
| Field | 1 | 810 | 295 | TIFT | C810 |
| Field | 1 | 811 | 19 | IL05-8515 | C811 |
| Field | 1 | 812 | 228 | 95AB12770 | C812 |
| Field | 1 | 813 | 379 | CORRAL | C813 |
| Field | 1 | 814 | 952 | M2-2 | C814 |
| Field | 1 | 815 | 194 | ANDREW | C815 |
| Field | 1 | 816 | 454 | PERDEBERG | C816 |
| Field | 1 | 817 | 165 | MN08130 | C817 |
| Field | 1 | 818 | 69 | X8826-1 | C818 |
| Field | 1 | 819 | 835 | IA111003 | C819 |
| Field | 1 | 820 | 610 | QUALITY\|PI289587 | C820 |
| Field | 1 | 821 | 323 | TX07CS1402 | C821 |
| Field | 1 | 822 | 584 | ONOHOJSKIJ_A-547 | C822 |
| Field | 1 | 823 | 332 | HORIZON270 | C823 |
| Field | 1 | 824 | 611 | CIAV6218 | C824 |
| Field | 1 | 825 | 460 | CIAV5389 | C825 |
| Field | 2 | 826 | 200 | EXCEL | C826 |
| Field | 2 | 827 | 228 | 95AB12770 | C827 |
| Field | 2 | 828 | 611 | CIAV6218 | C828 |
| Field | 2 | 829 | 295 | TIFT | C829 |
| Field | 2 | 830 | 19 | IL05-8515 | C830 |
| Field | 2 | 831 | 454 | PERDEBERG | C831 |
| Field | 2 | 832 | 194 | ANDREW | C832 |
| Field | 2 | 834 | 316 | TX02U7479 | C834 |
| Field | 2 | 835 | 835 | IA111003 | C835 |
| Field | 2 | 836 | 206 | WOODBURN | C836 |

| Location | Block | Plot | GID | T3Oat_Name[1] | RNASeq_Samplename[2] |
|---|---|---|---|---|---|
| Field | 2 | 837 | 610 | QUALITY\|PI289587 | C837 |
| Field | 2 | 838 | 952 | M2-2 | C838 |
| Field | 2 | 839 | 584 | ONOHOJSKIJ_A-547 | C839 |
| Field | 2 | 840 | 165 | MN08130 | C840 |
| Field | 2 | 841 | 332 | HORIZON270 | C841 |
| Field | 2 | 842 | 428 | PI344841 | C842 |
| Field | 2 | 845 | 69 | X8826-1 | C845 |
| Field | 2 | 846 | 124 | SHERWOOD | C846 |
| Field | 2 | 847 | 303 | FL0238BSB-22 | C847 |
| Field | 2 | 848 | 323 | TX07CS1402 | C848 |
| Field | 2 | 849 | 379 | CORRAL | C849 |
| Field | 2 | 850 | 460 | CIAV5389 | C850 |

[1]T3Oat_Name= line names were consistent with that on T3/Oat (https://triticeaetoolbox.org/oat/)
[2]RNASeq_Samplename= run name of 3' RNASeq samples
